# Integrated in silico and in vitro approaches identify a HSP90 drug vulnerability axis in t(7;12) infant leukemia stem-like cells

**DOI:** 10.1101/2025.09.06.674620

**Authors:** Ylenia Cicirò, Denise Ragusa, Emily M. Johnson, Ayona Johns, Sina Kanannejad, Remisha Gurung, Macarena Oporto Espuelas, Chun-Wai Suen, Gabriel Torregrosa-Cortés, Alice Giustacchini, Sabrina Tosi, Cristina Pina

**Author notes:** Address for correspondence: Cristina Pina, Sanquin Research, Plesmanlaan 125, 1066CX Amsterdam, The Netherlands. YC, DR and EMJ contributed equally to this work. DR and ST are co-corresponding authors.

## Abstract

The t(7;12) translocation is a chromosomal rearrangement characteristic of infant Acute Myeloid Leukemia (AML). It arises *in utero* and results in ectopic overexpression of homeobox gene *MNX1*. Using a 3-dimensional (3D) model of blood development, we recently showed that t(7;12)-AML originates at the endothelial-to-hematopoietic transition, explaining its characteristic gene expression signature. Herein, we employ that signature to interrogate the transcriptional profiles of hundreds of human cell lines against the GDSC database of drug sensitivities to identify candidate drugs against t(7;12)-AML. We employ a cell line in which we engineered t(7;12) and systematically test the candidate drugs by cell surface phenotype and clonogenic assays. Importantly, we identify HSP90 inhibitor SNX.2112 as a potential therapeutic agent against t(7;12)-AML. SNX.2112 selectively eliminates colony-initiating leukemia progenitors *in vitro* and decreases *MNX1* expression, effects recapitulated by other HSP90 inhibitors. SNX.2112 acts at least partly through destabilisation of STAT5 signalling. Critically, SNX.2112-differential signatures uniquely map to progenitors with hemato-endothelial characteristics in t(7;12)-AML patient blasts, suggesting targeting of leukemia-initiating cells. Combinatorial treatment with chemotherapeutic agents indicates synergy, suggesting SNX.2112 potential as a targeted and cytotoxicity-sparing therapeutic approach. Overall, we successfully use an integrated computational and multi-model experimental approach to identify a drug vulnerability of t(7;12)-AML.

**Key points:**

1. Classifier-based in silico drug screening identifies vulnerabilities of t(7;12)-infant leukemia
2. 2D and 3D models of t(7;12)-leukemia match HSP90 inhibition cellular and molecular responses to candidate leukemia stem cells in t(7;12) patient analysis.

## Introduction

Infant Acute Myeloid Leukemia (infAML) with t(7;12)(q36;p13) is exclusive to children under 2, encompassing 2-28% of infAML cases^1,2^. It is classified as high-risk in multiple trials; it associates with high relapse and low survival, but responds well to hematopoietic stem cell transplantation (HSCT)^2,3^. In absence of HSCT, therapeutic options remain limited. It was recently shown that t(7;12)-AML originates *in utero*^4^, matching our and others’ observations that t(7;12) and downstream ectopic *MNX1* expression are unable to transform adult cells^5–7^. *MNX1*-overexpression could initiate leukemia in fetal liver (FL) but not in bone marrow (BM) cells^7^. Complementarily, using a 3D-gastruloid model of embryonic blood development (hemogenic gastruloid, haemGx), we proposed that *MNX1*-overexpressing AML originates at the endothelial-to-hematopoietic transition (EHT) to emergence of erythroid-myeloid progenitors (EMP)^8^. Ectopic activation of *MNX1*, the hallmark of t(7;12), results from *ETV6* enhancer hijacking^9,10^.

We and others published engineered models of t(7;12), displaying *MNX1*-overexpression and capturing molecular signatures observed in patients^5,11^. In our model, t(7;12)(q36;p13) was engineered in K562 cells, superseding aspects of their BCR-ABL-driven oncogenic program to: (1) modify cellular composition and colony output and match the erythroid block and early myeloid bias of t(7;12)-AML; and (2) faithfully recapitulate transcriptional signatures separating t(7;12)-AML from healthy progenitors and other infAML forms^5^.

Herein, by interrogating a unique t(7;12) transcriptional signature against a large database of drug sensitivities, and testing candidate compounds for selective activity against K562-t(7;12) cells, we identified HSP90 inhibition as a putative vulnerability of t(7;12)-driven transformation. HSP90 inhibition down-regulated t(7;12)-associated genes with destabilized STAT5A signaling, and impacted hemato-endothelial signatures associated with embryonic blood specification, suggesting putative targeting of persisting embryonic progenitors at the root of t(7;12) infAML initiation. Importantly, signatures of hemato-endothelial affiliation and HSP90 responsiveness could be retrieved from t(7;12) patients, highlighting the translational potential of the findings.

## Methods

### Computational screening

Drug sensitivity (IC50) and expression data (RMA) for cell lines of the Genomics of Drug Sensitivity in Cancer (GDSC) project were downloaded from GDSC1000. Resistant and sensitive classifications of cell lines was retrieved as determined by dose-fitting curves by GDSC. Receiver-operating characteristic (ROC) analysis was performed in R using pROC to calculate area under the curve (AUC). Limma (v3.64.1 in R) was used for differential expression between sensitive *vs.* resistant lines.

### Cell culture

K562-t(7;12)/control lines were cultured as described before^5^. Hemogenic gastruloids with *MNX1*-overexpression (MNX1-oe-haemGx) were assembled as previously described^8^.

### Pharmacological treatments

Treatments were performed using pre-dissolved drugs from DiscoverProbe L1052-APE library (APExBIO*) vs*. 0.1% DMSO (control). Combination treatments were performed at IC50, ½IC50 or ¼IC50. Proliferation was determined by MTT (Cayman Chemical) as per manufacturer’s instructions. Absorbance was quantified at 570nm using CLARIOstar Plate Reader (BioTek). IC50 was calculated with GraphPad Prism. Drug synergy was analyzed by Highest Single Agent (HSA) using SynergyFinder+^12^.

### Colony-forming assays (CFC)

K562-t(7;12)/control were plated onto StemMACS™ HSC-CFU (Miltenyi Biotec), 500 cells/well (6-well plate). Disassembled MNX1-oe-haemGx were plated onto methylcellulose HSC007 (R&D Systems), 2×10^5^ cells/35mm-dish. Plates were incubated at 37°C, 5%CO_2_ for 10 days before scoring.

### Flow cytometry and cell sorting

Staining was performed in flow cytometry buffer (PBS/2%FBS/0.5mM EDTA) for 30min on ice (antibodies in **Table 1**). Cell tracking was performed in 5µM Tag-it Violet™ Proliferation and Cell Tracking Dye (Biolegend) for 30min at 37°C. Apoptosis was quantified using AnnexinV (Biolegend #640905) in AnnexinV Binding Buffer (1:20). Analysis was performed on ACEA Novocyte (Agilent). Cell sorting was performed using Beckman Coulter CytoFlex SRT (488nm 50mW, 633nm 100mW, 405nm 90mW, and 561nm 30mW), set with 100μm nozzle at 15PSI, using 4-way purity. Cells were gated from FSC-A/SSC-A and FSC-A/FSC-H plus SSC-A/SSC-H for doublet exclusion.

**Table 1.** List of antibodies.

| Antibody | Manufacturer | Catalogue number | Application and dilution |
| --- | --- | --- | --- |
| PE anti-human-CD117/c-Kit | Biolegend | 313203 | Flow cytometry – 1:100 |
| APC/Cyanine7 anti-human-CD24 | Biolegend | 311131 | Flow cytometry – 1:200 |
| BrilliantViolet785 anti-mouse-CD117/c-Kit | Biolegend | 105841 | Flow cytometry – 1:100 |
| BrilliantViolet510 anti-mouse-CD24 | Biolegend | 101831 | Flow cytometry – 1:100 |
| Anti-STAT5a antibody [E289] | Abcam | ab32043 | Western blot – 1:1000 |
| Anti-phospho-STAT5 Y694 | Abcam | ab32364 | Western blot – 1:1000 |
| GAPDH Monoclonal Antibody | Invitrogen | ZG003 | Western blot – 1:10000 |
| IRDye® 680RD Goat Anti-Rabbit IgG | LICOR | 926-68071 | Western blot – 1:5000 |
| IRDye® 800CW Donkey Anti-Mouse IgG | LICOR | 926-32212 | Western blot – 1:5000 |

### Western blot

Proteins were prepared by direct lysis in 1X Laemmli Buffer and separated by 12% SDS-PAGE and transferred on nitrocellulose membranes (Amersham). Membranes were blocked in 5%BSA for 1hr, blotted with primary antibodies diluted in TBST/BSA and secondary antibodies (**Table 1**), and visualized using Odyssey DLx (LICOR).

### Imaging

Cytation5 Cell Imaging Multi-Mode Reader (Biotek) was used for imaging using bright-field and FITC channels.

### RNA-sequencing

Total RNA was extracted using Monarch Total RNA Miniprep Kit (NEB). Sequencing was performed on NovaSeqX Plus PE150, at 50M paired-end reads/sample. Tophat2 with Bowtie2 were used to map to *Homo sapiens* genome GRCh39 and annotated using GENCODE V39. Aligned reads were quality-filtered (q30) using samtools^13^ and quantified using htseq^14^. Differential expression was computed using DESeq2^15^ in R (v1.48.0). Gene Ontology analysis was performed in EnrichR^16^. Gene Set Enrichment Analysis (GSEA) was performed on GSEA v4.2.3^17^, with parameters: 10000 permutations, gene set enrichment, weighted Signal2Noise. GDC TARGET-AML data (STAR counts and clinical information) was downloaded from https://ocg.cancer.gov/programs/target. GSVA was performed in R (v.2.2.1).

### Single-cell RNA-sequencing (scRNA-seq)

MNX1-oe-haemGx (*vs.* empty vector, EV) were collected at 216h, sorted as CD41^+^ and/or CD45^+^, and deposited into 96-well plate for Smart-Seq2 at 500kb and depth of 101 bases. Reads QC and trimming, alignment, and multimodality distributions were performed as described^8^.

ScRNA-seq data of t(7;12) patient sample and healthy control (ethics approval NHS REC16/LO/1290) were obtained as count matrices from the pediatric AML dataset in^18^, performed on 5’ 10x Genomics Chromium (Chromium Next GEM Single Cell 5′ Reagent Kits v2 Dual Index) and CITE-seq/Feature Barcode. Alignment and quantification used Cell Ranger (v7.1.0) against reference genome GRCh38. Ambient RNA and ADT contamination were removed using CellBender on raw Cell Ranger counts, applying false positive rates (FPR) 0.01 (RNA) and 0.1 (ADT).

Datasets were processed in Seurat (v.5.3.0) in R^19^, loaded and normalized individually using SCTransform and integrated by IntegrateData (SCT normalisation). Integrated data were analyzed with: PCA; shared nearest neighbor (SNN) clustering with 0.5 resolution; UMAP. Cluster markers were computed using FindAllMarkers by differential expression using Wilcoxon Rank Sum test. AUC enrichment scores were computed using AUCell (v.1.30.1).

### Statistical analysis

Experiments were performed at least in triplicates, unless specified otherwise. Statistical analysis was performed in R (V4.1.3) or GraphPad Prism 8.0.

## Results

### Correlating t(7;12) and drug sensitivity transcriptional signatures identifies candidate compounds with inferred activity against t(7;12)-AML

We previously defined t(7;12)-unique signatures from TARGET bulk RNA-seq to distinguish t(7;12) from other forms of pediatric AML^5^, and hypothesized that this signature could configure specific drug vulnerabilities. We correlated t(7;12) signature with drug responsiveness by probing 970 cell lines in the GDSC database with available gene expression and drug sensitivity data^20^. We selected the top 17 GSEA leading edge-genes enriched for t(7;12) signatures in the engineered K562-t(7;12) line^5^, and used them as a classifier to discern between sensitive and resistant cell lines in respect of the 264 drugs in GDSC. The cumulative capacity of the 17-gene signature to correctly call responsiveness to each drug was determined by Receiver Operating Characteristic (ROC) analysis (**Figure 1A**), making use of binary sensitive *vs.* resistant line classification in GDSC. The analysis yielded 33 compounds with AUC≥0.7, representing good correspondence between drug-dependent and t(7;12)-specific gene expression (**Figure 1B, File S1**). Reassuringly, when probed using signatures from other common infAML subtypes^5^, we found minimal overlap of drug responsiveness with t(7;12) (∼17%), indicating classifier specificity (**Figure 1S1A**, **File S1**). Seven of the 33 compounds selected for putative t(7;12)-AML responsiveness were available for downstream analysis in our laboratory in the DiscoverProbe library (APExBIO). We also included 2 drugs targeting HDAC6 (ACY1215, ACY241), substituting for CAY10603 retrieved by ROC. For all drugs selected, the t(7;12)-17-gene signature configured a reproducible pattern of differential gene expression between resistant and sensitive GDSC lines (**Figure 1C**).

**Figure 1.**
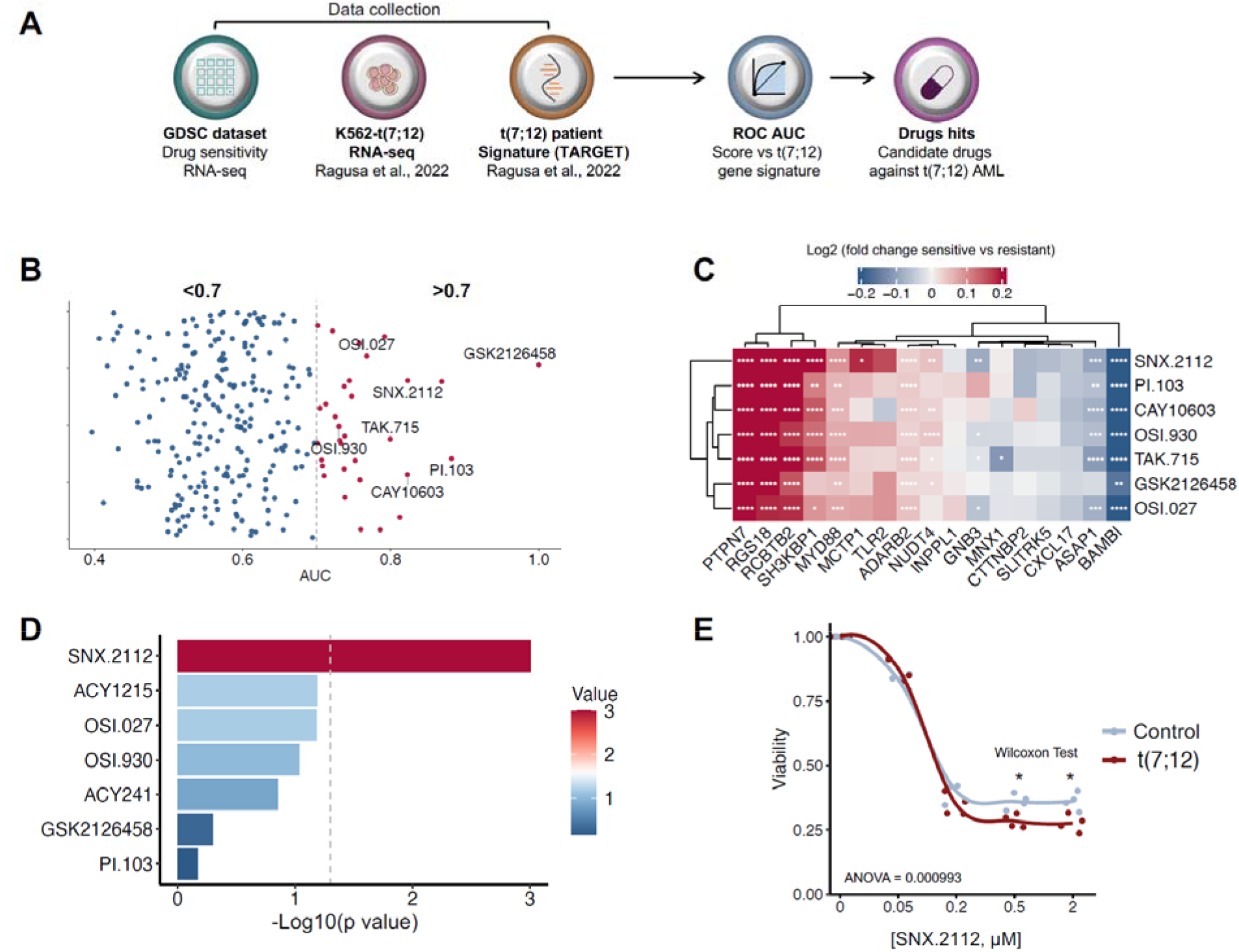
Interrogation of drug sensitivity databases with t(7;12) transcriptional signature identifies candidate pathways and compounds with inferred activity against t(7;12)-AML. **A.** Computational screening workflow. **B.** Cloud plot of all of drugs with enriched association with 17-gene t(7;12) signature filtered for area under the curve (AUC) computed by Receiver Operating Characteristic (ROC) analysis at >0.7. Random values are assigned to y axis to improve visibility of individual scattered data points. AUC values >0.7 were deemed significant. Highlighted names indicate the drugs tested *in vitro*. **C.** Heatmap representing the difference in gene expression between sensitive and resistant lines from GDSC for each drug, expressed as log2 fold change computed on differentially expressed genes determined by *limma*. Statistically significant genes by FDR > 0.05 are shown by asterisks as: p value ≤ 0.05 (*), 0.001 (**), 0.0001 (***), and 0.00001 (****). Hierarchical clustering by Euclidean distances. **D.** Summary of p values (expressed as - log10) from two-way analysis of variance (ANOVA) of IC50/dose curves across all drugs, highlighting compound-specific proliferation differences between t(7;12) and control K562 cells. The grey dashed line indicates the threshold of significance (-log10(0.05)). **E.** MTT dose response curve showing the cell viability of t(7;12) and control (transient Cas9-transfection only) K562 cells as a function of varying the SNX.2112 concentration. K562 cells were seeded at a concentration of 10000 cells and 25000 cells per well (U-bottom 96-well plate) for control and t(7;12) cells, respectively, for comparable densities at endpoint. N=3 with 2 technical duplicate measurements. Statistical significance was determined using the Wilcoxon rank-sum test with significant threshold set at p ≤ 0.05, and two-way ANOVA with interaction. P value ≤ 0.05 (*), 0.001 (**), 0.0001 (***), and 0.00001 (****).

We tested drug activity *in vitro* using the K562-t(7;12) line we previously engineered^5^, and tested differential responsiveness against the control line (K562-control). While all drugs showed dose-dependent proliferation inhibition by MTT metabolic assay (**Figure 1S1B**), only SNX.2112 showed significantly enhanced proliferation inhibition in K562-t(7;12) (**Figure 1D-E**). OSI.930 and OSI.027 showed a favourable, but non-significant trend towards enhanced inhibition in t(7;12); surprisingly, the 2 HDAC6 inhibitors had the opposite trend (**Figure 1S1B**). PI3K inhibitors GSK2126458 and PI.103 had no selective advantage. TAK.175 was excluded due to failure to reach 50% inhibition in both cell lines at 20mM, suggesting toxicity and low efficiency.

Taken together, *in silico* screening of gene expression profiles of drug responsiveness against a t(7;12)-specific signature identified a small number of compounds of interest for selective targeting of t(7;12)-infAML. Amongst these, HSP90 inhibitor SNX.2112 showed the greatest potential.

### SNX.2112 reduces colony-forming activity in t(7;12)-K562 cells

The transformed nature of K562 cells can confound proliferation assays, independently of the engineered t(7;12) translocation. We previously showed that the presence of t(7;12) alters K562 biology and specifically modifies colony-formation in methylcellulose-based assays, with increased proportion of “sparse” colonies of undifferentiated appearance^5^ (**Figure 2A**). DMSO (vehicle) treatment preserves colony-formation including sparse morphology (**Figure 2S1A**), confirming the assay utility in this setting, measuring culture-initiating progenitor activity as leukemogenic potential.

**Figure 2.**
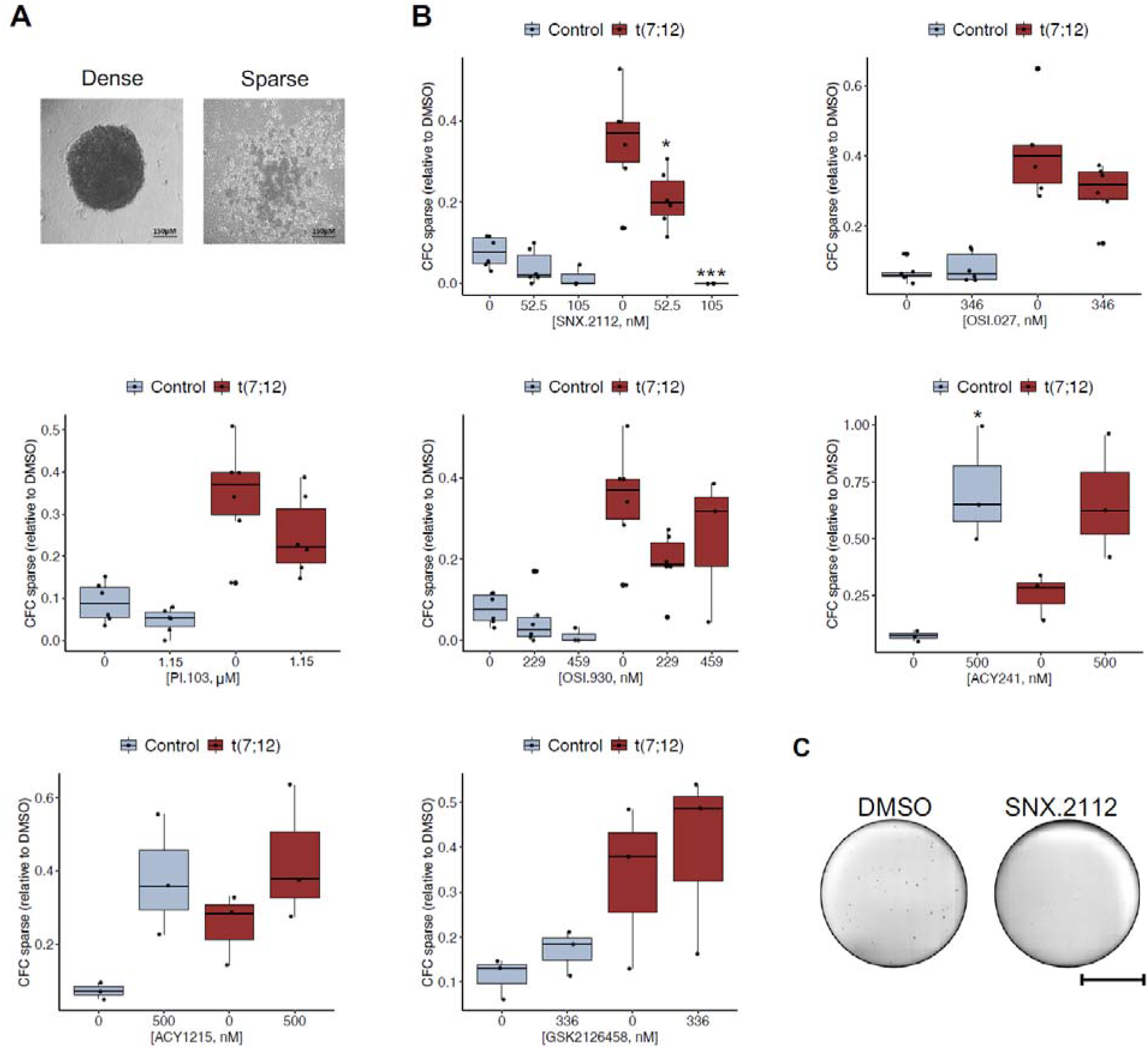
HSP90 inhibitor SNX.2112 reduces t(7;12) clonogenic activity. **A.** Representative images of dense and sparse colony types, in control (left) and t(7;12) K562 cells (right). **B.** Quantification of sparse colony-forming cell (CFC) upon treatment relative to the DMSO of the selected drugs in t(7;12) and control cells. Box plots with individual data points; statistical analysis was performed as a 2-way ANOVA with Tukey’s post-hoc multiple testing at significant p value ≤ 0.05. P value ≤ 0.05 (*), 0.001 (**), 0.0001 (***), and 0.00001 (****). **C.** Representative CFC images of t(7;12) K562 cells treated with DMSO (left) as control and with SNX.2112 drug (right). Scale bar, 10000μm.

Quantification of colonies upon treatment with the candidate compounds showed that SNX.2112 reduced total (**Figure 2S1B**) and sparse colony-formation (**Figure 2B**) in a dose-dependent manner, specifically in K562-t(7;12). Despite PI.103 and OSI.027 trends in total colony reduction (**Figure 2S1B-C**), these did not extend to sparse colonies (**Figure 2B**), highlighting SNX.2112’s selective activity against t(7;12)-carrying cells (**Figure 2B-C**). Interestingly, HDAC6 inhibitor ACY241 (trend in ACY1215), achieved the opposite effect of increasing sparse colonies, selectively in K562-control (**Figure 2B**). The paradoxical gain in sparse colony-formation is reversed in K562-t(7;12), in line with the expected direction of activity of our computational screen (**Figure 1S1B**). Taken together, analysis of colony-forming activity confirms the putative selectivity of SNX.2112 against t(7;12) leukemia cells, including targeting of candidate leukemia progenitors that sustain leukemia formation and are critical for therapeutic targeting.

### SNX.2112 targets colony-initiating Kit^+^CD24^+^ cells through proliferative block

To define K562-t(7;12) candidate leukemia progenitors, we employed surface markers Kit and CD24, previously shown by us to be expanded in engineered K562-t(7;12) cells^5^. Given t(7;12)-AML Kit-positivity^6,21^ and the specific association of CD24 with oncogenic *MNX1* expression^22^, we tested if these markers captured the K562-t(7;12) characteristic sparse colony morphology. We sorted K562-t(7;12) and K562-control on the 4 combinations of Kit/CD24 surface expression (**Figure 3S1A**), and quantified colony morphologies in methylcellulose-assays. Sparse colony frequency significantly associated with Kit-positivity (±CD24), exclusively in K562-t(7;12) (**Figure 3S1B**), attesting to differential biological properties of Kit^+^, including Kit^+^CD24^+^, cells in K562-t(7;12) cultures.

**Figure 3.**
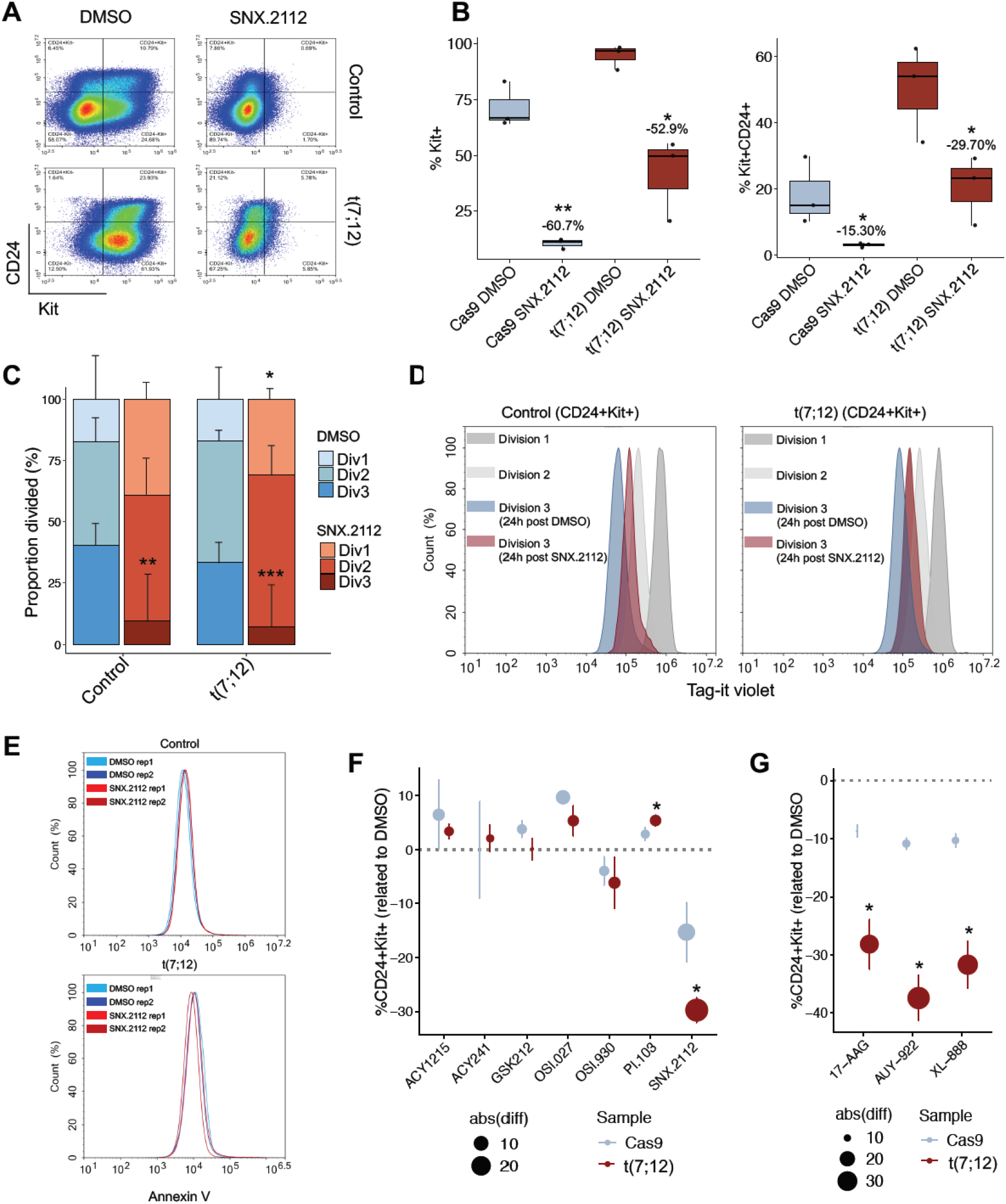
SNX.2112 selectively reduces Kit^+^ cells in t(7;12) K562 cells. **A.** Representative flow cytometry analysis of SNX.2112 effects on t(7;12) and control K562 cells. Data plotted as Kit^+/-^CD24^+/-^ gated on live singlets. **B.** Percentage of Kit^+^ (left) and Kit^+^CD24^+^ (right) cells from flow cytometry analysis of control- and t(7;12)-K562 treated with SNX.2112, compared to DMSO treatment. Percentage labels show the reduction as mean difference between %Kit^+^ or %Kit^+^CD24^+^ between DMSO *vs.* SNX.2112 treatment. Statistical significance calculated by t-test between DMSO and treatment; p value ≤ 0.05 (*), 0.001 (**), 0.0001 (***), and 0.00001 (****). **C.** Quantification of divisional tracking showing the proportion of cells undergoing division(s) under treatment with SNX.2112 or control (DMSO). Div1, div2, and div3 indicate the number of divisions. Statistical significance was determined by t-test with significant threshold set at p value ≤ 0.05; p value ≤ 0.05 (*), 0.001 (**), 0.0001 (***), and 0.00001 (****). **D.** Representative flow cytometry plots of Tag-it Violet intensities at each division comparing SNX.2112 and DMSO treatments in control- (left) and t(7;12)-K562 (right). **E.** Representative apoptosis analysis of control and K562-t(7;12) cells treated with DMSO or SNX.2112. Cells were stained with Annexin V and analysed after 5 h of treatment. Rep1 and rep2 indicate replicates. **F-G.** Reduction of Kit^+^CD24^+^ cells in control and K562-t(7;12) cells treated with other drugs from the ROC analysis (**F.**) or other HSP90 inhibitors (**G.**), calculated as difference between %Kit^+^CD24^+^ in DMSO control *vs.* treatment. Size represents the absolute value of difference in %; error bars show +/- standard deviation. Statistical significance calculated by t-test between cell lines; p value ≤ 0.05 (*), 0.001 (**), 0.0001 (***), and 0.00001 (****).

Consistent with the colony-formation response, SNX.2112 treatment significantly reduced the proportion of Kit^+^ and Kit^+^CD24^+^ cells, with a significantly steeper reduction in K562-t(7;12) where the Kit^+^CD24^+^ fraction is expanded (**Figure 3A-B**).

Selective loss of Kit^+^CD24^+^ cells by SNX.2112 was underpinned by reduction in proliferation with a higher inhibitory effect in K562-t(7;12) (**Figure 3C-D**), but no increase in apoptosis (**Figure 3E**). The other candidate drugs failed to significantly reduce the proportion of Kit^+^ and/or Kit^+^CD24^+^ cells in K562-t(7;12), with only OSI.930 showing a similar, albeit non-significant trend (**Figure 3F, 3S1C**). Importantly, 3 other HSP90 inhibitors (XL-888, AUY-922, and 17-AAG) ablated Kit^+^CD24^+^ cells (**Figure 3G, 3S1D**), highlighting selectivity of HSP90 inhibitory activity against the t(7;12)-associated Kit^+^CD24^+^ population. To further understand unique biological properties of candidate t(7;12), Kit^+^CD24^+^ leukemia progenitor cells, we performed RNA-seq comparing this fraction in K562-t(7;12) to K562- control (**Figure 3S1E** and **File S2**). We confirmed enrichment of t(7;12)-specific gene signatures, including *MNX1*-overexpression, in K562-t(7;12) Kit^+^CD24^+^ cells (**Figure 3S1E-F**). K562-t(7;12) Kit^+^CD24^+^ showed higher expression of developmental hemato-endothelial genes (*PROCR*, *CD44, RUNX1T1*, *ERG*, *ICAM2*, *LMO2*) and of early myelo-erythroid markers (*EPOR*, *SPI1*), while differentiated myeloid markers *(CSF3R, CD33*, *CD84*) were downregulated (**Figure 3S1E**). Cell type enrichment analysis supported the association of Kit^+^CD24^+^ with hemato-endothelial identities, reminiscent of the origin of *MNX1*-driven leukemia transformation at EHT we recently proposed^8^ (**Figure 3S1G**). Neuronal and pancreatic associations of K562-t(7;12) Kit^+^CD24^+^ cells reflect MNX1’s developmental role^23,24^.

Overall, SNX.2112 could target Kit^+^CD24^+^ progenitors, which are expanded by t(7;12) and associate with enhanced embryonic hemato-endothelial characteristics, reminiscent of candidate cell-of-origin of t(7;12)-infAML.

### SNX.2112 affects hemato-endothelial specification and colony-formation in *MNX1*-overexpressing hemogenic gastruloids

We previously proposed a hemato-endothelial temporal window of susceptibility to *MNX1*-overexpression (*MNX1-*oe), as a surrogate of t(7;12)-AML, using 3D haemGx^8^. We tested SNX.2112 activity in the same haemGx model, probing its effects on early developmental progenitors from which the leukemia may initiate; we focused on the 144-hour timepoint we pinpointed as susceptible to *MNX1-*oe^8^. *MNX1*-oe-haemGx were treated with SNX.2112 in 3D culture between 120-144 hours, capturing the MNX1-modified EHT. *MNX1*-oe-haemGx cells were also tested in serially-replated colonies dissociated from the 144-hour time-point to represent effects of *in vitro* transformation (**Figure 4A**).

**Figure 4.**
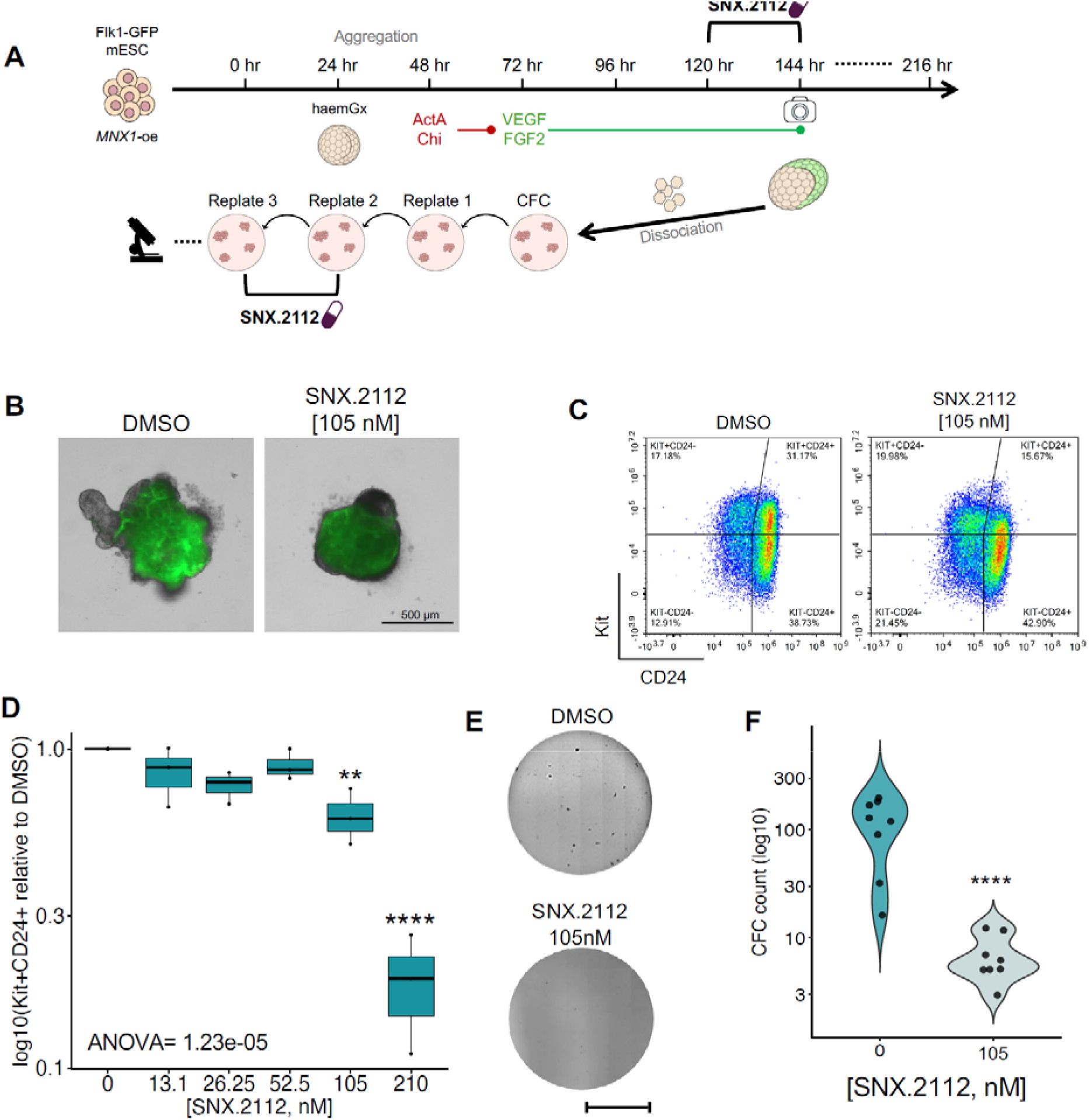
*MNX1*-overexpressing mouse hemogenic gastruloids (haemGx) lose transformation phenotype with SNX.2112 treatment. **A.** Schematic of treatment regimen in haemGx, depicting the timeline of growth and growth factors involved. Treatment of 3D haemGx or 2D haemGx-derived CFC is shown at 120 hr and replate 2, respectively. The icons indicate the readout methods: imaging at 144 hr for the treated haemGx, and microscope investigation and CFC counting for CFC. Haem-Gx-MNX1-oe were treated between 120 and 144h of culturing. Haem-Gx-MNX1-oe-derived CFC were treated at second replating for ∼10 days before scoring. **B.** Representative images of *Flk1-*GFP fluorescence of *MNX1*-oe-haemGx treated at day 5 (120 hr) of hemogenic specification with SNX.2112 or DMSO as control; haemGx were visualised 24 hr later. Treatment with selected dose of 105 nM is represented. Statistically significance was assessed via two-tailed t-test with a significance threshold of p value ≤ 0.05. **C.** Representative flow cytometry analysis of SNX.2112 effects on the haemGx model. Data plotted as Kit^+/-^CD24^+/-^. Cells are gated on live singlets. **D.** Quantification of %Kit^+^CD24^+^ cells upon treatment with escalating doses of SNX.2112 drug in the haemGx model. Values are plotted as log10 compared to the DMSO. Statistical analysis shown as a 2-way ANOVA with Tukey’s post-hoc multiple testing at significant p value ≤ 0.05; p value ≤ 0.05 (*), 0.001 (**), 0.0001 (***), and 0.00001 (****). **E.** Colony formation from serially-replated MNX1-oe cells obtained from 144 hr haemGx and analysed at third plating upon treatment with DMSO (control, up) or SNX.2112 (down) at a concentration of 105 nM, all in 0.1% DMSO. Scale bar, 10000μm. **F.** Quantification of the colony-forming capacity of the haemGx treated as in **(E.)** Independent replicates are shown. Statistically significance was assessed via two-tailed t-test with a significance threshold of p value ≤ 0.05; p value ≤ 0.05 (*), 0.001 (**), 0.0001 (***), and 0.00001 (****).

SNX.2112 treatment reduced endothelial *Flk1/KDR* (GFP reporter) signal in *MNX1-*oe-haemGx in a dose-dependent manner (**Figure 4B, 4S1A**), suggestive of endothelial program perturbation. Flow cytometry analysis of SNX.2112-treated *MNX1-*oe-haemGx revealed dose-dependent loss of Kit^+^CD24^+^ (**Figure 4C-D**), recapitulating observations in K562. In contrast, *MNX1-*oe-haemGx treated with OSI.930, a candidate drug that displayed mild reduction of Kit^+^CD24^+^ in K562 (**Figure 3S1C**), showed reduced GFP (*Flk1/KDR*) only at high doses (**Figure 4S1B-C**), and a mild albeit significant reduction of Kit^+^CD24^+^, which is not dose-dependent (**Figure 4S1D**). SNX.2112 treatment of serially-replated *MNX1-*oe-haemGx cells resulted in near-complete loss of colony-forming potential (**Figure 4E-F**), an effect not observed with OSI.930 (**Figure 4S1E-F**).

We corroborated these observations by scRNA-seq analysis of *MNX1*-oe-haemGx cells *vs.* control (**Figure 4S2A, File S3**). *MNX1*-oe-enriched cluster 3 (**Figure 4S2B**) had a significantly enhanced representation of t(7;12)-AML (**Figure 4S2C**) and SNX.2112-response signatures (**Figure 4S2D-F**). Conversely, no specific association with *MNX1*-oe cells was found for OSI.930-responsive signature (**Figure 4S2G-H**). Overall, the data are consistent with an inhibitory effect of SNX.2112 on propagation of t(7;12) progenitor-like leukemia cells with hemato-endothelial features, using independent cell models.

### SNX.2112 targets t(7;12)-AML-specific signatures and modifies STAT5A signalling

Having established that SNX.2112 perturbs unique aspects of t(7;12)-AML biology, we interrogated downstream molecular programs underpinning these effects. We performed RNA-seq analysis of SNX.2112-treated K562-t(7;12) cells *vs.* DMSO unfractionated cultures (**Figure 5A, File S4**); attempts to retrieve the Kit^+^CD24^+^ sub-population were frustrated by low persistence in culture after SNX.2112 treatment.

**Figure 5.**
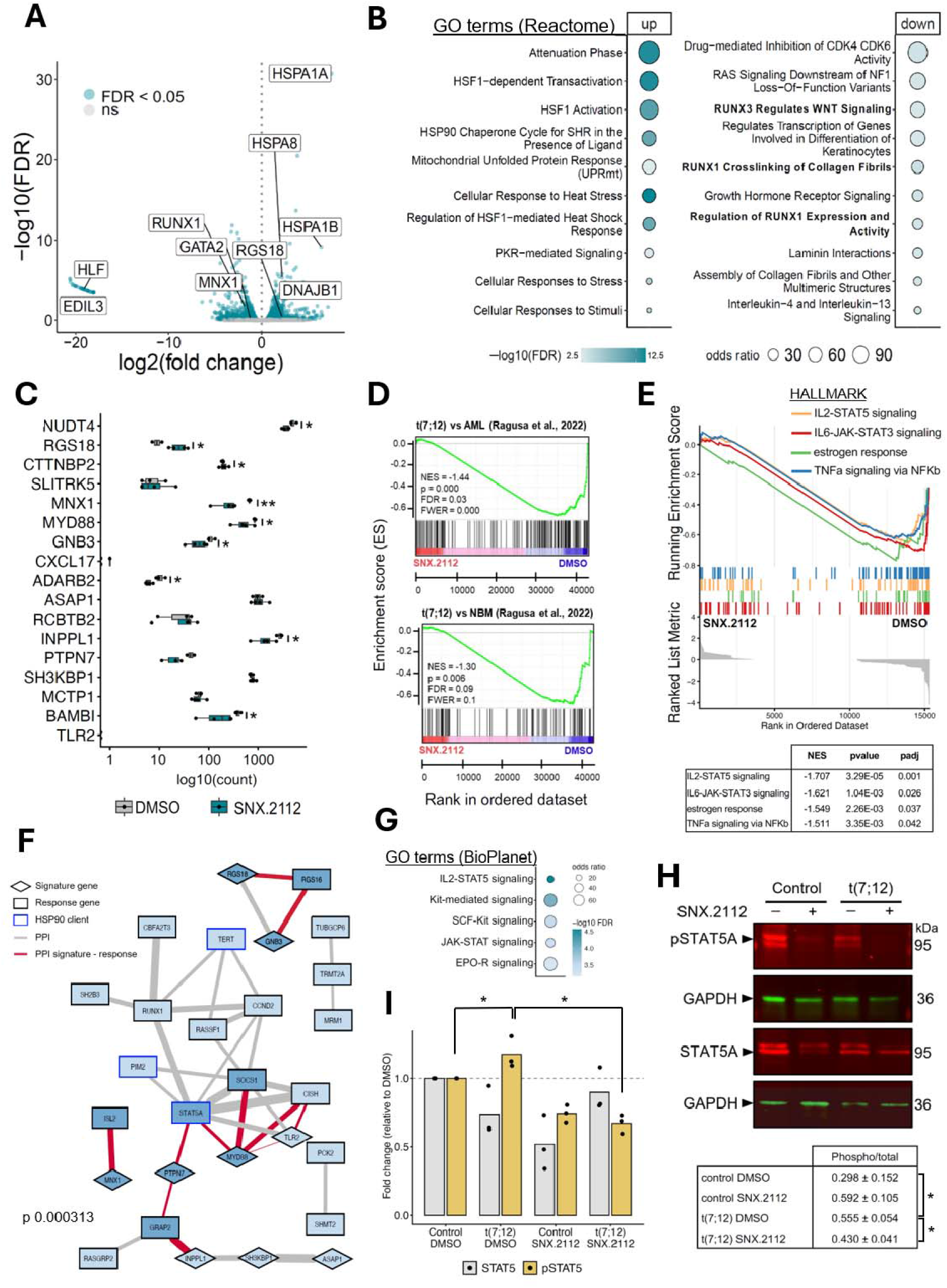
SNX.2112 treatment of t(7;12) K562 cells affects expression of signature genes, including *MNX1*. **A.** Volcano plot of RNA-seq analysis of K562-t(7;12) cells treated with SNX.2112 vs DMSO control cells; respectively 4 and 3 independent samples. Differentially-expressed genes (DEG) at FDR <5% are represented in turquoise. **B.** Gene Ontology (GO) term enrichment for differentially upregulated genes between SNX.2112-treated and DMSO-treated K562-t(7;12) cells, computed in EnrichR using the Reactome repository. **C.** Gene expression levels of the 17-gene signature used for ROC analysis in K562-t(7;12) cells treated with SNX.2112 or DMSO control. Statistical significance was determined using two-tailed t-test with significant threshold set at p value ≤ 0.05; p value ≤ 0.05 (*), 0.001 (**), 0.0001 (***), and 0.00001 (****). **D.** GSEA of the t(7;12) vs AML and t(7;12) vs NBM signatures^5^ against RNA-seq of K562-t(7;12) treated with SNX.2112 (red) or DMSO (blue). Normalized enrichment scores (NES) are deemed significant by p val<0.05 and FDR <0.1. **E.** GSEA of hallmark gene signatures against RNA-seq of K562-t(7;12) treated with SNX.2112 or DMSO. Normalized enrichment scores (NES) are deemed significant by p val<0.05 and FDR <0.1, and are reported in the table below. **F.** STRING analysis of known protein-protein interactions (PPI) combining the t(7;12) 17-gene signature with the SNX.2112 response signature. PPIs (shown as edges) were set to default STRING settings to include textmining, experiments, databases, co⍰ZIexpression, neighborhood, gene fusion, and co⍰ZIoccurrence (confidence threshold ≥ 0.4); genes with no interactions were removed from the network. Edge thickness represents confidence scores for PPI evidence strength. Node shapes represent genes from t(7;12) 17-gene signature (diamond) and SNX.2112 response gene (rectangle shape). A blue border indicates a known HSP90 client protein. Nodes are filled in dark blue for significantly downregulated genes (by FDR < 0.05) in K562-t(7;12) treated with SNX.2112 that form connections between a t(7;12) gene and a response gene; light blue indicates other downregulated genes; white indicate statistically non-significant changes in expression. Red edges represent PPIs between a t(7;12) gene and a response gene that are statistically downregulated (FDR < 0.05). Grey edges represent PPIs of all query genes. **G.** Gene Ontology (GO) term enrichment for genes forming the PPI network in F., computed in EnrichR using the BioPlanet 2019 repository. **H.** Representative Western blot image of K562-control and K562-t(7;12) treated with SNX.2112 or DMSO (indicated by “-“), probed for phospho(p)STAT5A and total STAT5A, with GADPH as housekeeping control. Quantification of intensity by ratio of pSTAT5A/total STAT5A is shown in the table below, showing +/- standard deviation of N=3. Statistical significance was determined using t-test with significant threshold set at p value ≤ 0.05 (*). **I.** Quantification of Western blot analysis of phospho(p)STAT5A and total STAT5A by intensity, normalized to GAPDH, and represented as fold change compared to K562-control treated with DMSO. Statistical significance was determined using the t-test with significant threshold set at p value ≤ 0.05 (*).

Genes up-regulated by SNX.2112 included heat-shock proteins (e.g. *HSPA1A-B, HSPA8*), matching enrichment of gene ontologies (GO) representing heat-shock-related pathways and unfolded protein response (UPR), consistent with HSP90 inhibition (**Figure 5A-B**). Multiple GO capturing SNX.2112 down-regulated genes corresponded to RUNX1 regulatory activity (**Figure 5B**), encompassing down-regulation of *RUNX1* itself and other key hematopoietic regulators (*GATA2*, *HLF*), and endothelial genes (e.g. *EDIL3*) (**Figure 5A**). This pattern is again reminiscent of the putative embryonic EHT origin of t(7;12)-AML^8^, seemingly directly perturbed by SNX.2112. Importantly, *MNX1* and the 17-gene signature employed for computational screening were down-regulated upon SNX.2112 treatment (**Figure 5A,C**). None of these genes were down-regulated upon OSI.930 treatment (**Figure 5S1A-B**), which fails to target t(7;12)-specific features and impacts generic MYC-driven oncogenic programs instead (**Figure 5S1C-D**). GSEA confirmed significant negative enrichment of t(7;12)-signatures^5^ in SNX.2112-treated K562-t(7;12) *vs.* DMSO (**Figure 5D**), but not with OSI.930 (**Figure 5S1D**). Transcriptional changes for both t(7;12) and SNX.2112-response signatures were limited in treated K562-control cells; reassuringly, HSP90-inhibitor XL888 elicited a transcriptional response comparable to SNX.2112 (**Figure 5S1E**).

Unsupervised inspection of hallmark gene-sets significantly depleted upon SNX.2112 treatment retrieved STAT signalling as a top target program (**Figure 5E**). Interestingly, STAT5 is a central node in a STRING protein-protein interaction (PPI) analysis to map putative links between the original 17-gene signature and SNX.2112 response signatures (**Figure 5F, 4S2D**). This captures multiple signature-response connections, including MNX1, and comprises STAT5A (itself, a HSP90 client), as well as RUNX1. In contrast, the OSI.930 response network showed minimal connectivity with the t(7;12)-signature, restricted to MYD88, and involving MYC (**Figure S41F**); no connectivity with MNX1 was inferred.

Interacting nodes between SNX.2112-response and t(7;12) signature (**Figure 5F**) captured JAK/STAT signalling targets (**Figure 5G**), suggesting differential STAT5 activation/response in K562-t(7;12) *vs.* K562-control. Western blot analysis of total and phospho(p)STAT5A showed that t(7;12)-carrying cells had higher baseline STAT5A activation as measured by phospho/total STAT5A ratio (**Figure 5H**). Conversely, pSTAT5A was specifically targeted by SNX.2112 treatment in t(7;12)-K562 cells, while treatment of K562-control preferentially reduced total STAT5 levels (**Figure 5H-I**).

Altogether, the data suggest that t(7;12)-specific leukemia programs are differentially responsive to SNX.2112 treatment, not only quantitatively in comparison with control cells, but qualitatively, through altered impact on STAT5 activation, a link between t(7;12) and SNX.2112-responsiveness signatures.

### Single-cell-RNA-seq analysis of t(7;12)-patient blasts maps SNX.2112 response to t(7;12)-enriched clusters

We asked whether SNX.2112-response signatures and differentially-targeted pathways in K562-t(7;12) are present in AML patients. We initially interrogated TARGET^25^ bulk RNA-seq data from infAML patients. The SNX.2112-response signature (**Figure 4S2D**) was preferentially enriched in t(7;12) patients compared to normal bone marrow (NBM) or to common subtypes KMT2A-rearrangements and inv(16) (**Figure 6A**). OSI.930 response was depleted in all infAML *vs.* NBM (**Figure 6S1A**). Using the SNX.2112-response STRING network we constructed earlier (**Figure 5F**), SNX.2112-response genes upregulated in t(7;12) *vs.* other infAML and/or NBM (**Figure 6B**) showed significant coverage of the network (**Figure 6C**, pink shading), suggesting putative translatable axes in t(7;12)-AML; nodes included MNX1, RUNX1 and STAT5A.

**Figure 6.**
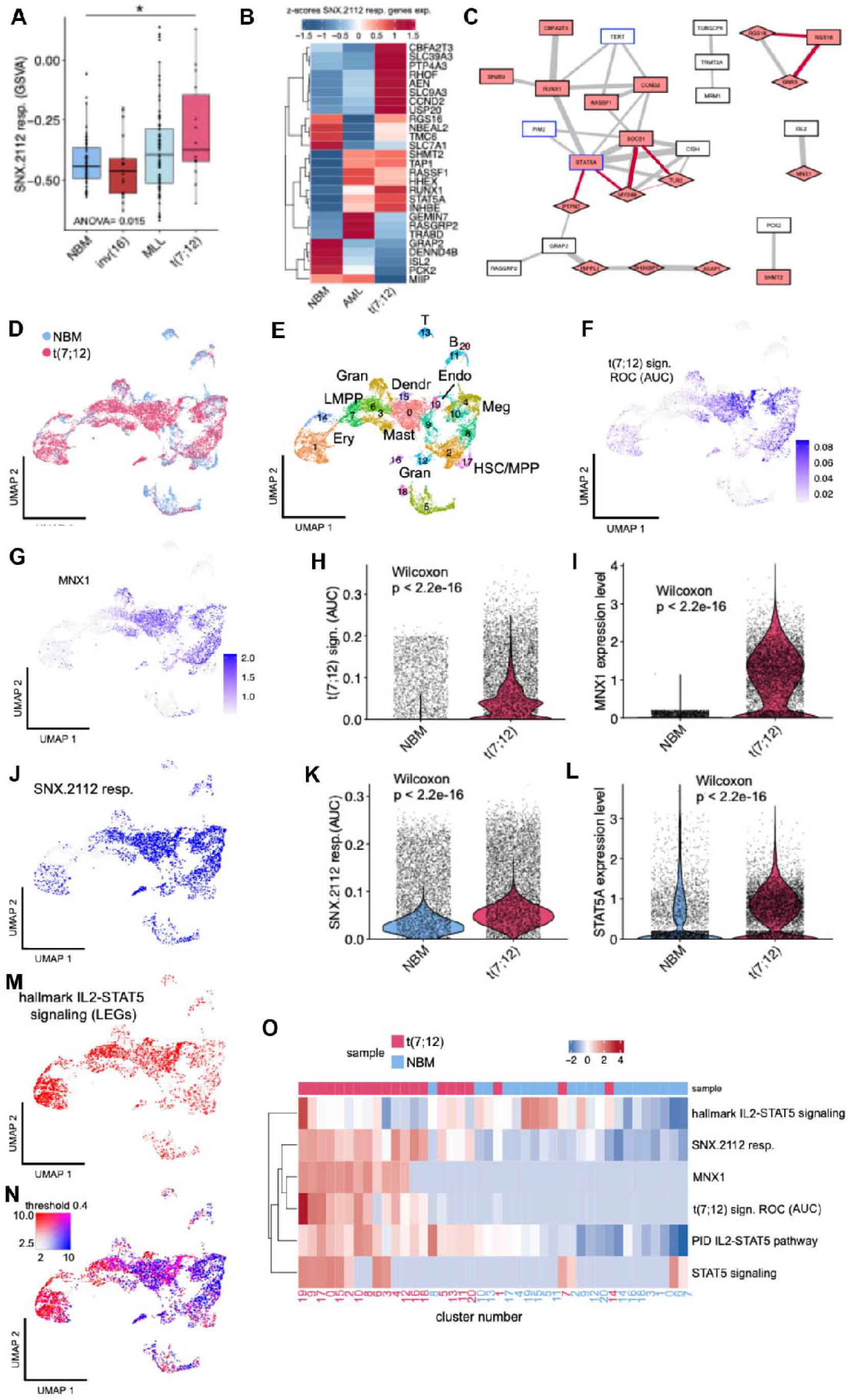
scRNA-seq of t(7;12)-AML patients reveals enrichment in SNX.2112 drug responsiveness signature. **A.** GSVA enrichment analysis of SNX.2112 response signature in infant AML bulk-RNA seq from TARGET by subtype [inv(16), MLL/KMT2A, and t(7;12)] and normal bone marrow (NBM). Statistical testing was calculated by ANOVA and post-hoc pair-wise significance between classes by Tukey’s test; p value ≤ 0.05 (*), 0.001 (**), 0.0001 (***), and 0.00001 (****). **B.** Heatmap of expression levels of SNX.2112 response genes in t(7;12), infant AML [inv(16) and MLL/KMT2A] as Z-scores, and normal bone marrow (NBM) from bulk RNA-seq data from TARGET. **C.** STRING analysis of known protein-protein interactions (PPI) combining the t(7;12) 17-gene signature with the SNX.2112 response signature (as described in Figure 5F), highlighting statistically significant upregulated (by Wilcox test, p val < 0.05) genes in RNA-seq of t(7;12) patients *vs.* other AML, or vs. normal bone marrow (NBM). **D-E.** UMAP embedding of scRNA-seq of integrated dataset of CD34+ cells from a t(7;12)-AML patient and paediatric normal bone marrow (NBM), coloured by **(D)** by cell identity of the original sample, and **(E)** cluster number identified by and labelled by cell identity based on differentially expressed genes. **F.** UMAP showing the enrichment value of the 17-gene t(7;12) signature used for ROC analysis, in AUC units calculated by AUCell, from lowest enrichment expression values in white to highest in dark blue. **G.** UMAP showing the expression of MNX1. **H.** Violin plot comparing AUC values for t(7;12) signature from ROC analysis between normal bone marrow (NBM) and t(7;12)-AML patient. Statistical significance is determined by Wilcoxon test. **I.** Violin plot comparing the expression of MNX1 between normal bone marrow (NBM) and t(7;12)-AML patient. Statistical significance is determined by Wilcoxon test. **J.** Co-expression analysis of SNX.2112 response (J., in AUC values) and hallmark IL2-STAT5 signaling (M., in AUC values) mapped onto UMAP, and shown as overlap of blue and red as magenta (N., co-expression threshold 0.4 shown in matrix). **K.** Violin plot comparing AUC values for SNX.2112 response between normal bone marrow (NBM) and t(7;12)-AML patient. Statistical significance is determined by Wilcoxon test. **L.** Violin plot comparing the expression of STAT5A between normal bone marrow (NBM) and t(7;12)-AML patient. Statistical significance is determined by Wilcoxon test. **M-N.** Co-expression analysis of SNX.2112 response (J., in AUC values) and hallmark IL2-STAT5 signaling (M., in AUC values) mapped onto UMAP (see J.). **O.** Heatmap of pseudo-bulk mean values of expression (MNX1 and STAT5A) or AUC enrichment values (response / pathway signatures) shown as Z-scores across clusters divided by normal bone marrow (NBM, in blue) and t(7;12)-AML patient (in pink). Hierarchical clustering by complete method on Euclidean distances.

We further dissected SNX.2112 responsiveness using scRNA-seq data of enriched CD34^+^ cells from a t(7;12) patient *vs*. NBM (**Figure 6D-E**). We annotated clusters based on lineage-affiliated genes (**File S5**), confirming broad progenitor representation (**Figure 6E**). We mapped t(7;12) cell proportion in each cluster (**Figure 6S1B**), and identified clusters 0, 1, 6, 8, 16 and 19 as uniquely enriched in (7;12)-AML blasts (**Figure 6S1C**). Clusters 0,1, and 6 captured erythroid and mast cell progenitors (**Figure 6E**), consistent with lineage affiliations^8^ and erythroid differentiation block^26^ of *MNX1*-overexpression. Cluster 19 configured an endothelial identity, matching analysis of *MNX1*-oe-haemGx^8^. The t(7;12)-17-gene signature used for computational drug screening (**Figure 6F**), and *MNX1* expression (**Figure 6G**), map to endothelial (19), mast/MEP (0) and HSC (8)-enriched clusters (**Figure 6S1C**); they are overall enriched amongst t(7;12) cells (**Figure 6H-I**). Significantly, the SNX.2112-response signature was also enriched in t(7;12)-CD34^+^ patient blasts, with broad recapitulation of t(7;12) cluster distribution (**Figure 6J-K**). No/mild enrichments were observed for the OSI.930-response signature (**Figure 6S1D**). Given the putative relevance of STAT5 signalling for t(7;12) vulnerability to SNX.2112, we next inspected the representation of STAT5-responsive signatures from treated K562-t(7;12) in patient scRNA-seq data. Unsurprisingly, STAT5 signatures, like STAT5A itself, were enriched in the MEP/erythroid/mast cell progenitor clusters, putatively reflecting KIT signalling (**Figure 6L**). Overlap between STAT5-signalling and SNX.2112-response signatures (**Figure 6M**, magenta) was particularly marked in endothelial-like cluster 19, specifically in the t(7;12)-patient cells (**Figure 6O**), reminiscent of the putative origin of the leukemia^8^. Broadly, we observed a close coincidence between t(7;12), SNX.2112 and STAT5 signatures in patient blasts in all clusters but the lymphoid-affiliated 5, 7, 11, 13, 14, and 20, where STAT5 reflects immune response pathways but no MNX1/t(7;12)-program is activated (**Figure 6O**). Overall, the data suggests that STAT5-mediated vulnerabilities to SNX.2112 may be present in t(7;12) patients’ AML blasts. The findings reinforce the notion that these may constitute targetable pathways with specificity to t(7;12) leukemia, including at the level of candidate leukemia-originating cells.

### SNX.2112 synergises with chemotherapeutic agents against K562-t(7;12) progenitors

Finally, we tested SNX.2112 in combination with chemotherapeutic agents cytarabine and mitoxantrone, both used in infAML induction schemes^27,28^. We calculated synergistic effects using HSA, to answer whether combinatorial treatment is superior to single-agent for enhanced cell viability reduction at lower doses. Paired combinatorial testing of SNX.2112 with cytarabine (**Figure 7A-C**) or mitoxantrone (**Figure 7D-F**) using MTT, showed significant synergistic effects, including at the lowest doses of both paired drugs (**Figure 7A,D**), denoting potential for reduced therapeutic dosages. In contrast, no synergistic effects were observed between OSI.930 and the chemotherapeutic agents (**Figure 7S1A-C,D-F**), or between the chemotherapeutic agents themselves (**Figure 7S1G-I**). Inspection of gene-sets associated with response to cytarabine or mitoxantrone showed significant enrichment in SNX.2112-treated K562-t(7;12) (**Figure 7G**), more strongly for cytarabine signatures, with no significant enrichments upon OSI.930 treatment (**Figure 7S1J**).

**Figure 7.**
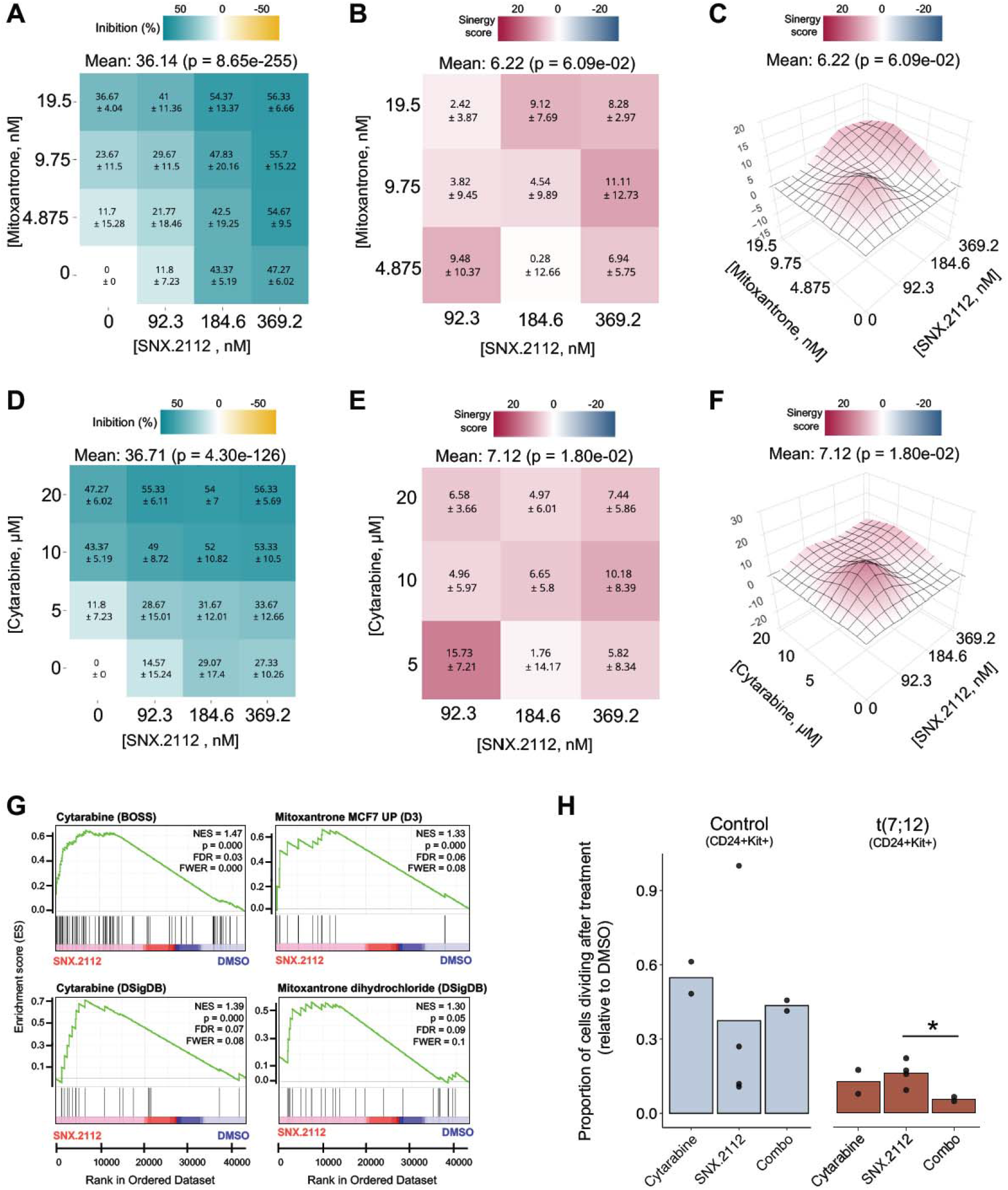
SNX.2112 synergises with chemotherapeutic agents to hinder proliferation of Kit^+^CD24^+^ of K562-t(7;12) cells. **A-D.** Dose response map of K562-t(7;12) treated with SNX.2112 in combination with mitoxantrone (**A.**) or with cytarabine (**D.**). Values of ¼ IC50, ½ IC50, and IC50 of all the drugs are indicated. p values at the whole dose matrix level are shown at the top of the matrixes, together with the global mean. Individual scores for each combination are shown together with the ± value of the mean for each replicates set. **B-E.** HSA synergy heatmaps showing the synergy score of K562-t(7;12) treated with SNX.2112 in combination with mitoxantrone (**B.**) or with cytarabine (**E.**). A score >1 was considered synergistic. Values of ¼ IC50, ½ IC50, and IC50 of all the drugs are indicated. p values at the whole dose matrix level are shown at the top of the heatmaps, together with the global mean. Individual scores for each combination are shown together with the ± value of the mean for each replicates set. **C-F.** 3D representation of the **B.** and **E.** HSA synergy maps, respectively. **G.** GSEA of the cytarabine (BOSS and DSigDB databases) and mitoxantrone (MCF7 UP(D3) and DSigDB databases) signatures against RNA-seq of K562-t(7;12) treated with SNX.2112 (red) or DMSO (blue). Normalized enrichment score (NES) are deemed significant by p val<0.05 and FDR<0.1. **H.** Proportion of Kit+CD24+ fraction undergoing cell divisions after treatment with SNX.2112 (92.3 nM) and cytarabine (5 mM), alone or in combination. K562-control cells are shown as light blue and K562-t(7;12) are represented in burgundy. Individual data points are shown and values are represented relative to DMSO treated control. Statistical significance is determined by Wilcoxon test and symbolized as: p value ≤ 0.05 (*), 0.001 (**), 0.0001 (***), and 0.00001 (****).

Importantly, the effect of combined SNX.2112 and cytarabine treatments significantly inhibited proliferation specifically within the Kit^+^CD24^+^ compartment, an effect exclusively present in K562-t(7;12) cells (**Figure 7H**). The autofluorescence of mitoxantrone precluded reliable analysis by flow cytometry.

Overall, the data indicate synergism between SNX.2112 and chemotherapeutic agents which extends to putative elimination of candidate t(7;12)-AML progenitors. Clinical potential is also highlighted by the low SNX.2112 toxicity observed against healthy human CD34+ cells in *in vitro* liquid cultures at doses active against t(7;12)-K562 cells (**Figure 7S1K**).

## Discussion

We harnessed characteristic gene expression programs of t(7;12)-AML to identify HSP90 inhibitor SNX.2112 as a candidate compound with distinct activity against this infAML subtype, including modulation of STAT5 signaling. SNX.2112 selectively hindered proliferation of clonogenic t(7;12)-expanded Kit^+^CD24^+^ cells and decreased expression of t(7;12)-defining genes, showing synergism with cytarabine. SNX.2112-response signatures were specifically enriched in t(7;12) patient transcriptomes, suggesting translational potential.

We identified SNX.2112 through leveraging of transcriptomic and pharmacological data from the largest database of drug sensitivities in cancer cell lines, GDSC^20,29–31^. We employed the binary ROC classifier system to evaluate discriminatory power of a t(7;12)-specific AML signature separating sensitive and resistant lines to each database drug. ROC is widely employed to evaluate clinical parameter accuracy; its standardised AUC values constitute intuitive power thresholds suitable for intrinsically-heterogenous datasets^32,33^. Wide GDSC representation of drug classes enables unbiased search generalizable for drug repurposing. While our ROC prediction is based on a collection of cancer cell lines not exclusively leukemia, its predictive power lies in the ability to probe expression patterns independently of tissue specificity. This was advantageous for t(7;12) signatures, which we previously showed to capture features of endothelium and neuronal/pancreatic programs^8^, and could therefore be probed across various cellular backgrounds.

We tested candidate compounds using our t(7;12)-engineered K562 line^5^. K562 are a multi-lineage progenitor line with characteristics of fetal hematopoiesis^35,36^, a developmental affiliation that enabled successful t(7;12) engineering and facilitates recapitulation of its unique haemato-endothelial properties^5^. We matched t(7;12) features to a Kit^+^CD24^+^ phenotype that was present in both K562^5^ and *MNX1*-oe-haemGx^8^ models^8^. CD24 is a direct transcriptional target of MNX1^22^; Kit is present in t(7;12) blasts^1,6,37^. *In vivo*, Kit^+^CD24^+^ marks a transition state in erythroid commitment of yolk sac erythro-myeloid progenitors^38^, coinciding with the developmental stage we previously defined as susceptible to *MNX1*-oe^8^.

Importantly, SNX.2112 may direct or extend its anti-proliferative activity to subsets of cells responsible for initiation/propagation of t(7;12)-AML, targeting pathways relevant to hemato-endothelial identity. SNX.2112 is a specific HSP90 inhibitor, reported to suppress solid tumor growth *in vitro* and *in vivo*^40,41^, with anti-angiogenic and anti-osteoclastogenic activities in myeloma^42^. It is a chaperone to multiple client proteins, including hematopoietic kinases (i.e. JAK2, FLT3)^43^, and HIF-1α, which regulates endothelial KDR/VEGFR-dependent signalling^44^. HSP90 also forms complexes with globins to assist folding^45^, and has been implicated in subcellular localization of nuclear proteins, including HSC and erythroid/megakaryocytic regulator MLLT3^46–48^. Accordingly, sub-populations of t(7;12)-AML blasts with hemato-endothelial and erythoid transcriptional characteristics also displayed SNX.2112 sensitivity signatures, highly enriched for STAT5 signaling. We demonstrated that SNX.2112 specifically targets active phosphorylated STAT5A in t(7;12), but not in control K562 cells, suggesting that the translocation alters AML vulnerabilities to HSP90 targeting. STAT5 phosphorylation can be directly assisted by HSP90^49^. The putative centrality of STAT5 to t(7;12)-defining protein-interaction networks, including upstream of RUNX1 a critical regulator of blood development essential for EHT^50^, links t(7;12) drug sensitivity to leukemia origin, suggesting potential to target leukemia stem-like cells. The fact that the molecular vulnerability features are detectable in scRNA-seq analysis of a t(7;12) patient, specifically so in clusters of cells with endothelial programs, supports this view. As a caveat, this the first report of scRNA-seq profiling of a t(7;12) AML. Additional samples of this rare disease will be needed to enable precise delineation of leukaemia hierarchies and specific mechanisms of drug susceptibility.

SNX.2112 (and pro-drug SNX.5422) is orally bioavailable with improved safety profile compared to earlier HSP90 inhibitors^51^, showing favourable phase I tolerability profiles in lung cancer^52^ and hematological malignancies^53^. Pan-HSP90 inhibitors have progressed slowly in entering clinical practice for toxicity and commercial reasons, despite promising reduction of solid and haematological tumour burden, alone or in combination^40,54,55^. The ongoing development of next-generation isoform-specific HSP90 inhibitors will strengthen translational applications^56^. In the case of SNX.2112, we observed good tolerability by healthy cultured human CD34+ cells at doses toxic for K562-t(7;12) cells, suggesting a translatable therapeutic window. Furthermore, we observed that other HSP90 inhibitors with distinct biochemical mechanisms of action produced similar specific effects against K562-t(7;12) cells, re-enforcing the likelihood of identifying an agent of clinical utility against t(7;12)-infAML within the HSP90 drug family.

InfAML remains challenging to target due to its rarity and difficulties of conducting clinical trials in this age group. Combining improved biological models with systematic screening of drug sensitivities against disease-specific transcriptional signatures can direct drug identification and functional testing, with the potential to target leukemia stem-like cells, improving outcomes of treatment and survival. Our integrative ROC analysis classification approach proved successful in streamlining identification of targetable molecular axes relevant to patient t(7;12)-AML. In principle, similar strategies can be harnessed for translational discovery in rare tumours and other rare diseases with limiting availability of patient material and experimental models.

## Acknowledgements

This research was funded by the Little Princess Trust through the Children’s Cancer and Leukaemia Group: CCLGA (CCLGA 2023 22 Pina). DR is the recipient of a European Hematology Association (EHA)-EMBL/EBI Computational Biology Training in Hematology (CBTH) award (CBTH39). AJ is funded by a Lady Tata Memorial Trust Scholarship (2022-2025). The CP lab receives funding from NC3Rs - National Centre for Replacement, Reduction and Refinement of Animals in Research (NC/Z500677/1) and the ERC (ERC Synergy Grant 2024 MakingBlood). The Authors are grateful to Jordi Garcia-Ojalvo for helpful discussions regarding the analysis of scRNA-seq data from MNX1-oe hemogenic gastruloids.

## Authorship Contributions

CP, ST conceived the projected. CP, DR, YC designed experiments. EMJ, YC, DR, AJ, RG, CWS performed experiments. MOE, SK, AG contributed critical datasets. DR, GTC, SK curated data. YC, DR, GTC, CP analyzed data. CP, DR, YC interpreted data. YC, DR represented data. CP, DR, YC wrote the manuscript. CP, DR, YC, ST edited the manuscript. CP supervised the work. All Authors read and agreed to the final version of the manuscript.

## Data availability statement

RNA-seq raw data for K562 drug treatments and sorted fractions is available at Array Express under accession codes E-MTAB-15526 and E-MTAB-15530. Single-cell RNA-seq for the haemGx-MNX1-oe is available at Array Express with accession code E-MTAB-12149. The results published here are partly based upon data generated by the Therapeutically Applicable Research to Generate Effective Treatments (TARGET) (https://ocg.cancer.gov/programs/target) initiative, of the Acute Myeloid Leukemia (AML) cohort GDC TARGET-AML. The data used for this analysis are available at https://portal.gdc.cancer.gov/projects and https://xenabrowser.net/. Drug sensitivity and gene expression data from GDSC is available at https://www.cancerrxgene.org/.

