## Supplementary Figures and Legends for "Integrated in silico and in vitro approaches identify a HSP90 drug vulnerability axis in t(7;12) infant leukemia stem-like cells"

[1]Sanquin Research, Landsteiner Laboratory, Amsterdam, The Netherlands; [2]Department of Biosciences and [3]Centre for Genome Engineering and Maintenance, College of Health, Medicine and Life Sciences, Brunel University of London, Uxbridge, United Kingdom; [4]Human Technopole, Milan, Italy; [5]Molecular and Cellular Immunology Section, UCL Great Ormond Street Institute of Child Health, Zayed Centre For Research into Rare Disease in Children, London, United Kingdom; [6]Department of Genetics, University of Cambridge, Cambridge, United Kingdom; [7]EMBL Barcelona, Barcelona, Spain; [8]Cancer Center Amsterdam, Amsterdam UMC, Amsterdam, The Netherlands; [9]Princess Máxima Centre for Pediatric Oncology, Utrecht, The Netherlands

Footnotes:

*YC, DR and EMJ contributed equally to this work.

### Supplementary Files

**File S1:** List of GDSC1 represented drugs used for the computational screen with indication of AUC. Targets and pathways of drugs with AUC >0.7 are indicated.

**File S2:** RNA-seq DEG in Kit$^+$CD24$^+$ cells from K562-t(7;12) vs K562-control.

**File S3:** RNA-seq DEG in K562-t(7;12) cells treated with SNX.2112 or OSI.930 vs DMSO.

**File S4:** scRNA-seq analysis of *MNX1*-oe *vs*. EV haemGx indicating DEG per cluster.

**File S5:** scRNA-seq analysis of t(7;12) CD34$^+$ cells *vs.* healthy pediatric BM indicating DEG per cluster.

### Supplementary Figures and Legends

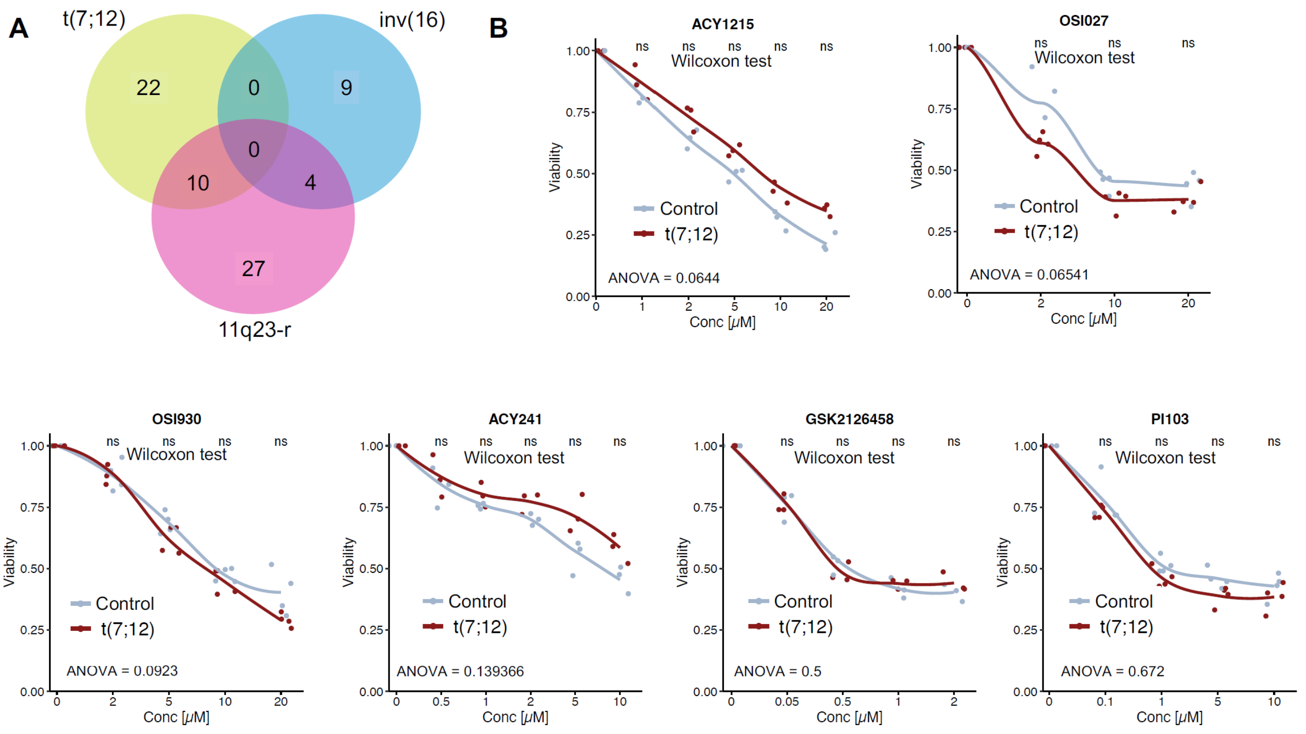


**<u>Figure 1 S1.</u> Screening of candidate compounds with putative activity against t(7;12).** **A.** Venn diagram showing the overlap of drug responsiveness between t(7;12) and other two forms of infAML: inversion 16 (inv(16)), and KMT2A/11q23 rearrangements (11q23-r). Responsiveness was attributed to drugs with AUC ≥0.7 of the binary classification value of their respective gene signature. **B.** MTT assay-based proliferation IC50 curves in t(7;12)- and control-K562 cells. N=3 with 2 technical duplicate measurements. Statistical significance was determined using the Wilcoxon rank-sum test with significant threshold set at p value ≤ 0.05, and two-way ANOVA with interaction; p value ≤ 0.05 (*), 0.001 (**), 0.0001 (***), and 0.00001 (****).

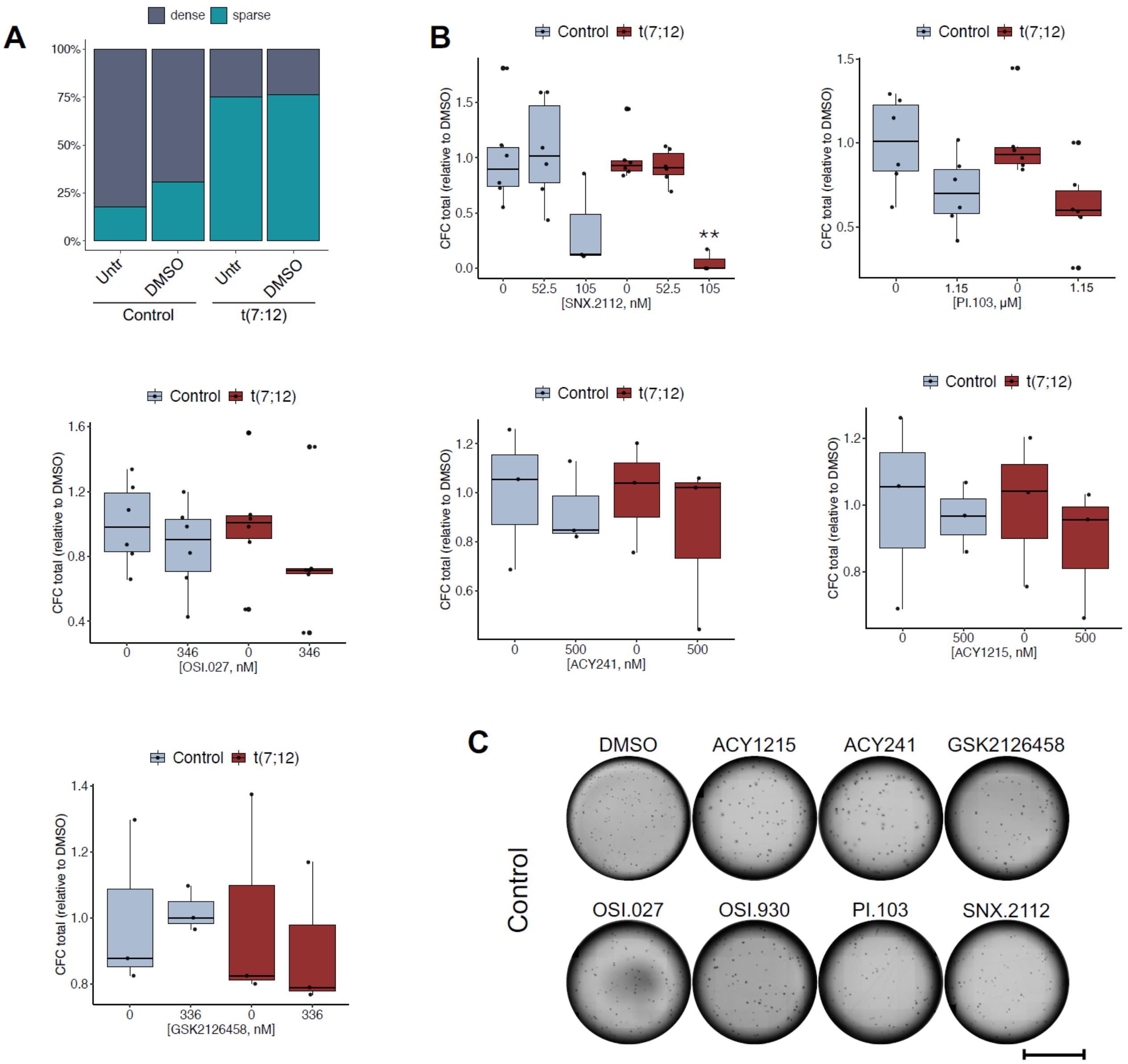


**<u>Figure 2 S1.</u> Candidate t(7;12) targeting drugs have differential effects on colony formation. A.** Validation of DMSO effects on colony formation; representative experiment. Untr, untreated cells. **B.** Quantification of total colonies upon treatments with escalating doses of the selected drugs. Box plots with individual data points. Statistical analysis by 2-way ANOVA with post-hoc Tukey's multiple testing, at significant p value ≤ 0.05, symbolized as p value ≤ 0.05 (*), 0.001 (**), 0.0001 (***), and 0.00001 (****). C. Overview of whole plates of CFC from control (up) or t(7,12) (down) treated cells. Scale bar, 10000µm.

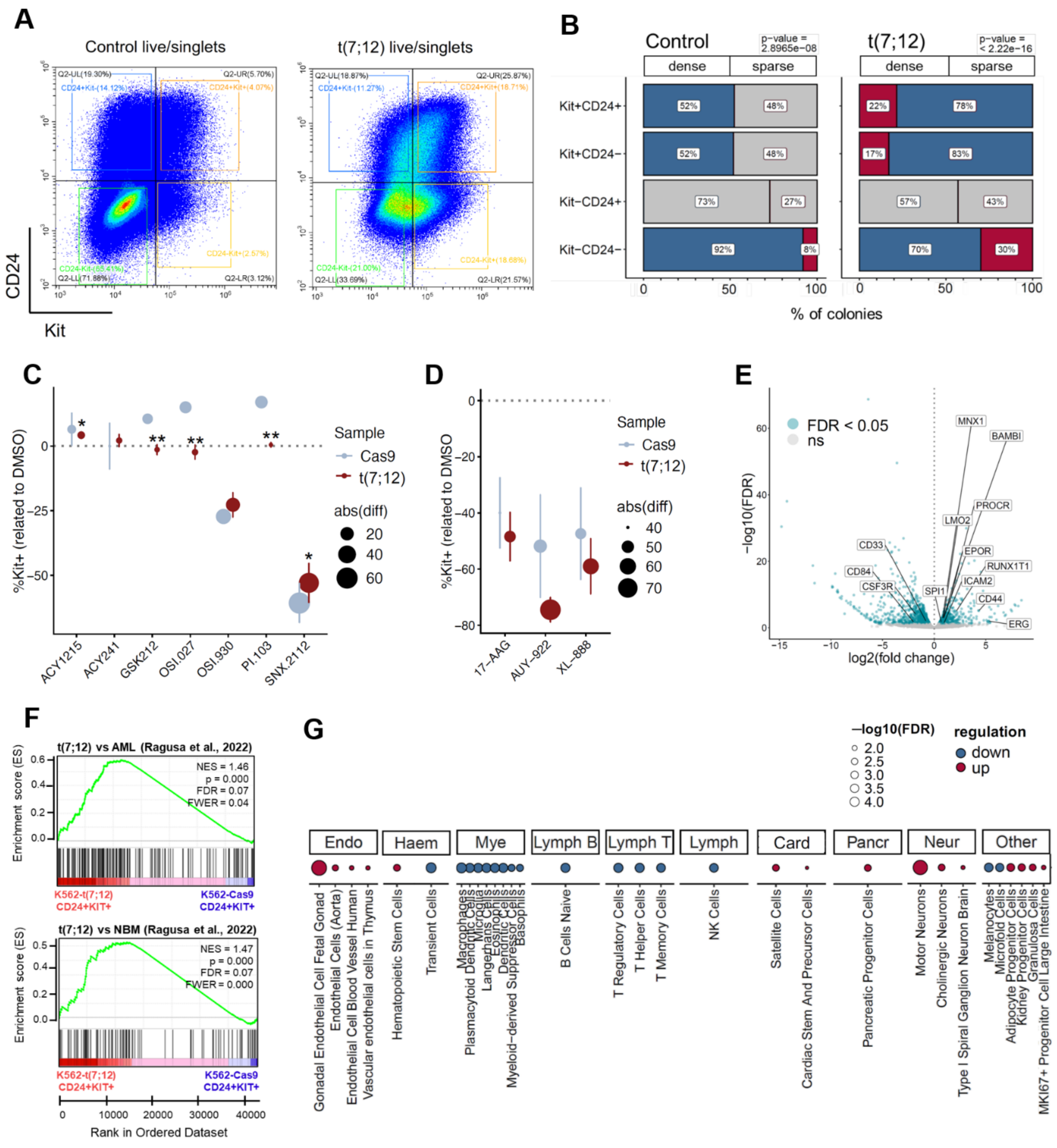


**<u>Figure 3 S1.</u> Sparse colony formation associates with Kit+ progenitors, specifically in t(7;12) K562 cells. A.** Gating strategy for sorting Kit and CD24 fractions in control and t(7;12) K562 cells. **B.** Proportion of CFC type (dense/sparse) in Kit+/-CD24+/- in control and t(7;12) translocated cells. Pearson residual as Chi-square test is shown. **C.** Reduction of Kit+ cells in control and t(7;12) K562 cells treated with other drugs from the ROC analysis, calculated as difference between %Kit+ in DMSO control vs. treatment. Size represents the absolute value of difference in %; error bars show +/- standard deviation. Statistical significance calculated by t-test between cell lines; p value ≤ 0.05 (*), 0.001 (**), 0.0001 (***), and 0.00001 (****). **D.** Reduction of Kit+ cells in control and t(7;12) K562 cells treated with other HSP90 inhibitors, calculated as difference between %Kit+ in DMSO control vs. treatment. Size represents the absolute value of difference in %; error bars show +/- standard deviation. Statistical significance calculated by t-test. **E.** Volcano plot showing the differentially expressed genes (DEG) at FDR < 5% identified in the RNA-seq analysis of Kit+CD24+ fraction of K562-t(7;12) compared to the K562-control. Fold change of expression is shown as –log10(FDR). Significance was set as FDR ≤ 0.05. N=3. Developmental haemato-endothelial genes and early myelo-erythroid markers are highlighted. **F.** GSEA of the t(7;12) vs AML and t(7;12) vs NBM signatures (Ragusa et al., 2022) against RNA-seq of Kit+CD24+ fraction of K562-t(7;12) vs. K562-control. **G.** Cell type enrichment analysis in Kit+CD24+ K562-t(7;12) cells vs. K562-control using representative cell type gene sets from Panglao DB 2021 database; bubble plot shows the type of regulation (up- or downregulated) and statistical significance (–log10(FDR), bubble size).

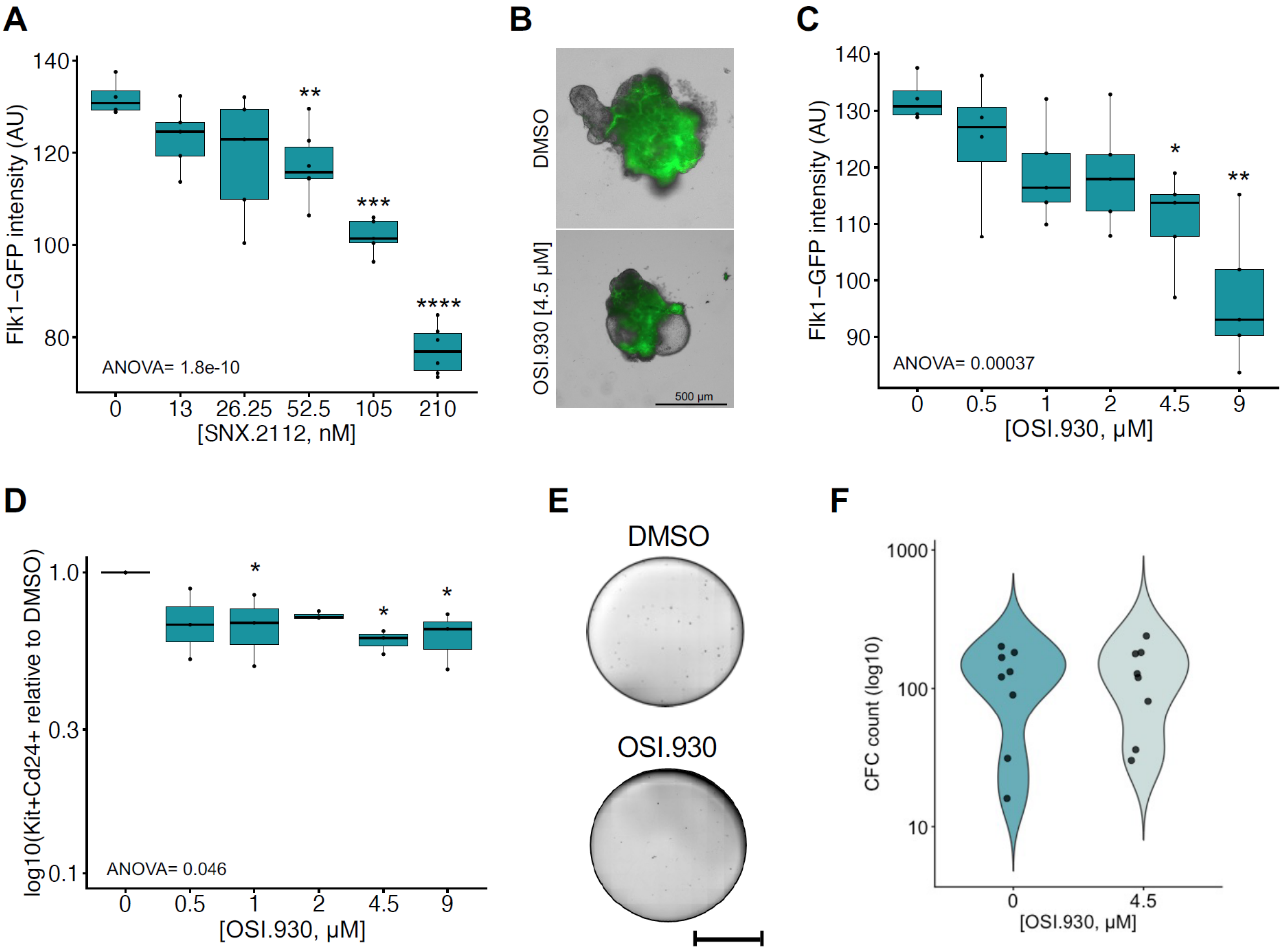


**<u>Figure 4 S1.</u> MNX1-overexpressing mouse haemogenic gastruloids (haemGx) lose transformation phenotype with OSI.930 treatment. A.** Quantification of Flk1- GFP signal (FITC channel, arbitrary fluorescence units AU) from images of MNX1-oe haemGx treated at escalating doses of SNX.2112. Statistical significance across doses was determined by ANOVA and post-hoc significance between doses by Tukey's test; p value ≤ 0.05 (*), 0.001 (**), 0.0001 (***), and 0.00001 (****). **B.** Representative images of Flk1-GFP fluorescence of MNX1-oe haemGx treated at day 5 (120 hr) of haemogenic specification with OSI.930 or DMSO; haemGx were visualized 24 hr later. Treatment with selected dose of 4.5 mM is represented. **C.** Quantification of Flk1-GFP signal (FITC channel, arbitrary fluorescence units AU) from images of MNX1-oe haemGx treated at escalating doses of OSI.930. Statistical significance across doses was determined by ANOVA and post-hoc significance between doses by Tukey's test; p value ≤ 0.05 (*), 0.001 (**), 0.0001 (***), and 0.00001 (****). **D.**

Quantification of % Kit+CD24+ 132 by flow cytometry upon treatment with escalating doses of OSI.930 drug in MNX1-oe haemGx. Values are plotted as log10 compared to the DMSO. Statistical significance across doses was determined by ANOVA and post-hoc significance between doses by Tukey's test; p value ≤ 0.05 (*), 0.001 (**), 0.0001 (***), and 0.00001 (****). **E.** Colony formation from serially-replated MNX1-oe cells obtained from 144 hr haemGx and analyzed at third plating upon treatment with DMSO (control) or OSI.930 at a concentration of 4.5 mM, all in 0.1% DMSO. Scale bar: 10000 μm. **F.** Quantification of colony-forming capacity as colony counts of MNX1-oe haemGx after treatment with OSI.930 or DMSO. Statistically significance was assessed via two-tailed t-test with a significance threshold of p value ≤ 0.05.

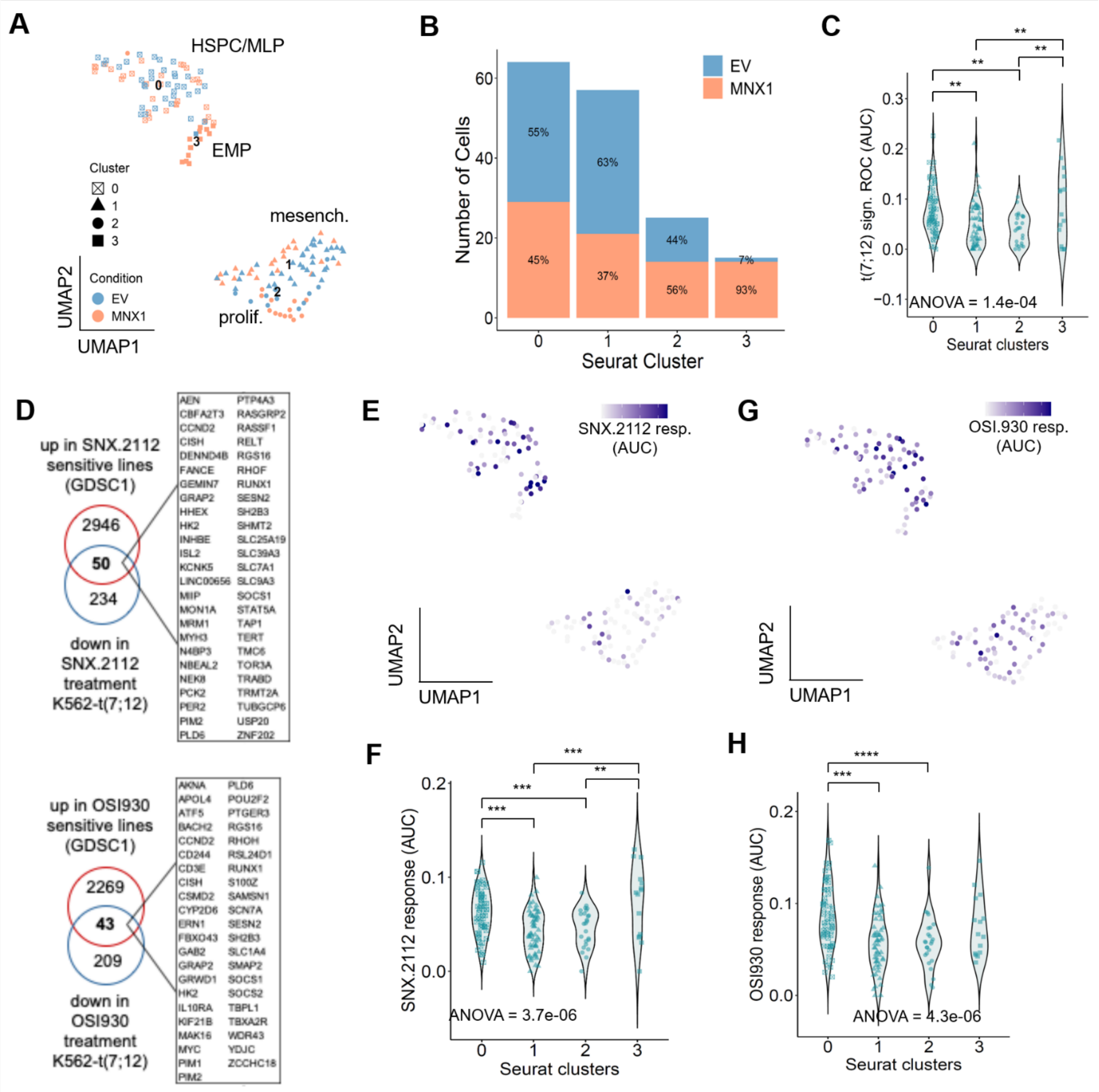


**Figure 4 S2. MNX1-overexpressing haemGx have unique or enriched cell types with over-representation of SNX.2112 responsiveness signatures. A.** UMAP embedding of scRNA-seq of integrated datasets of haemGx-MNX1-oe and haem-Gx empty vector (EV) at 216h sorted on CD45 and/or CD41, colour-coded by sample identity (MNX1 or EV), and coded by shape by cluster number labelled by cell type identity based on differentially expressed genes. **B.** Bar chart representing the proportion of cells populating each cluster by sample identity. **C.** Violin plot showing AUC values for t(7;12) signature from ROC analysis

by cluster number. Statistical testing was calculated by ANOVA across datapoints and post-hoc pair-wise significance between clusters by Tukey's test; p value ≤ 0.05 (*), 0.001 (**), 0.0001 (***), and 0.00001 (****). **D.** Venn diagram showing the intersect genes (complete list in zoomed box) used to generate the SNX.2112 (left) and OSI.930 (right) response gene sets, obtained by intersecting upregulated genes in sensitive lines to SNX.2112 or OSI.930 compared to resistant ones in the GDSC dataset (computed by limma, filtered by FDR < 0.05 and fold change > 0) and downregulated genes in K562-t(7;12) treated with SNX.2112 or OSI.930 compared to DMSO (computed by limma, filtered by FDR < 0.05 and fold change < 0). **E.** UMAP showing the enrichment value of SNX.2112 response in AUC units calculated by AUCell, from lowest enrichment expression values in white to highest in dark blue. **F.** Violin plot showing AUC values for SNX.2112 response by cluster number. Statistical testing was calculated by ANOVA across datapoints and post-hoc pair-wise significance between clusters by Tukey's test; p value ≤ 0.05 (*), 0.001 (**), 0.0001 (***), and 0.00001 (****). **G.** UMAP showing the enrichment value of OSI930 response in AUC units calculated by AUCell, from lowest enrichment expression values in white to highest in dark blue. **H.** Violin plot showing AUC values for OSI930 response by cluster number. Statistical testing was calculated by ANOVA across datapoints and post-hoc pair-wise significance between clusters by Tukey's test; p value ≤ 0.05 (*), 0.001 (**), 0.0001 (***), and 0.00001 (****).

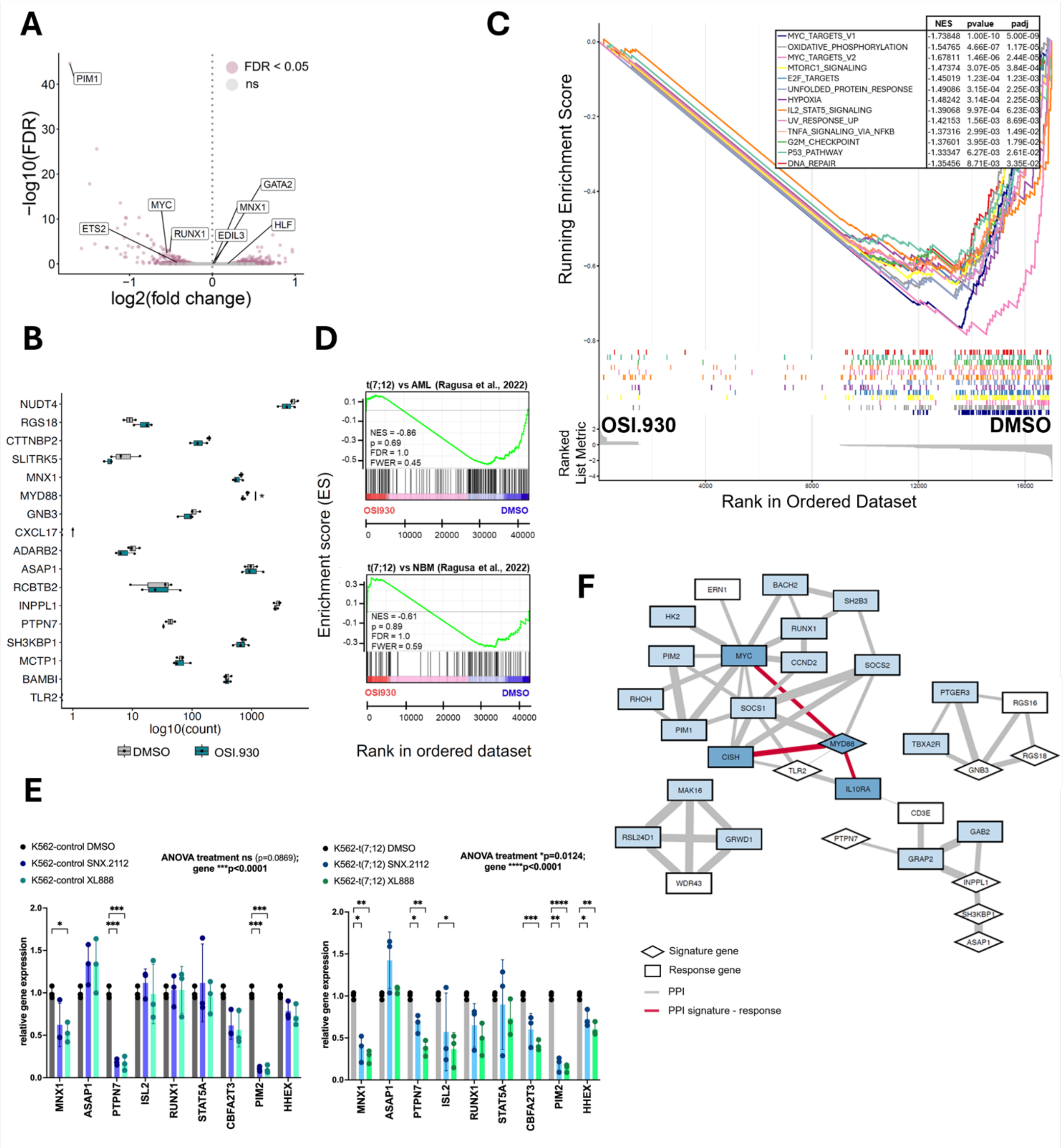


**<u>Figure 5 S1.</u> OSI.930 treatment of t(7;12) K562 cells does not impact expression of signature genes. A.** Volcano plot of RNA-seq analysis of K562-t(7;12) cells treated with OSI.930 vs DMSO control cells; respectively 4 and 3 independent samples. Differentially-expressed genes (DEG) at FDR < 0.05 are represented in pink colour. N=3 with 2 technical duplicate samples. **B.** Gene expression levels of the 17-gene signature used for ROC analysis in K562- t(7;12) cells treated with OSI.930 or DMSO control. Statistical significance was determined using the paired t-test with significant threshold set at p value $\leq 0.05$; p value $\leq 0.05$ (*), 0.001 (**), 0.0001 (***), and 0.00001 (****). **C.** GSEA of hallmark gene signatures against RNA-seq of K562-t(7;12) treated with OSI.930 or DMSO. Normalized enrichment scores (NES) are deemed significant by p val<0.05 and FDR <0.1, and are reported in the table on top. **D.** GSEA of the t(7;12) vs AML and t(7;12) vs NBM signatures (Ragusa et al., 2022) against RNA seq of K562-t(7;12) treated with OSI.930 (red) or DMSO (blue). **E.** Real-time quantitative (qPCR) analysis of selected SNX.2112- and t(7;12)- signature genes in K562-control and K562-t(7;12) treated with SNX.2112, XL888, and DMSO control. Relative gene expression fold change calculated by normalization to HPRT. Bars represent mean of 3 replicates and show individual data points; p values by ANOVA across datapoints; post-hoc statistical significance between specific variables by Tukey’s test and shown by brackets. **F.** STRING analysis of known protein-protein interactions (PPI) combining the t(7;12) 17-gene signature with the OSI.930 response signature. PPIs (shown as edges) were set to default STRING settings to include textmining, experiments, databases, co-expression, neighborhood, gene fusion, and co-occurrence; genes with no interactions were removed from the network. Node shapes represent genes from t(7;12) 17-gene signature (diamond) and OSI.930 response gene (rectangle shape). Nodes are filled in dark blue for significantly downregulated genes (by FDR < 0.05) in K562-t(7;12) treated with OSI.930 that form connections between a t(7;12) gene and a response gene; light blue indicates other downregulated genes; white indicate statistically non-significant changes in expression. Red edges represent PPIs between a t(7;12) gene and a response gene that are statistically downregulated (FDR < 0.05). Dashed edges represent PPIs of all query genes. Edge thickness represents confidence scores for PPI evidence strength.

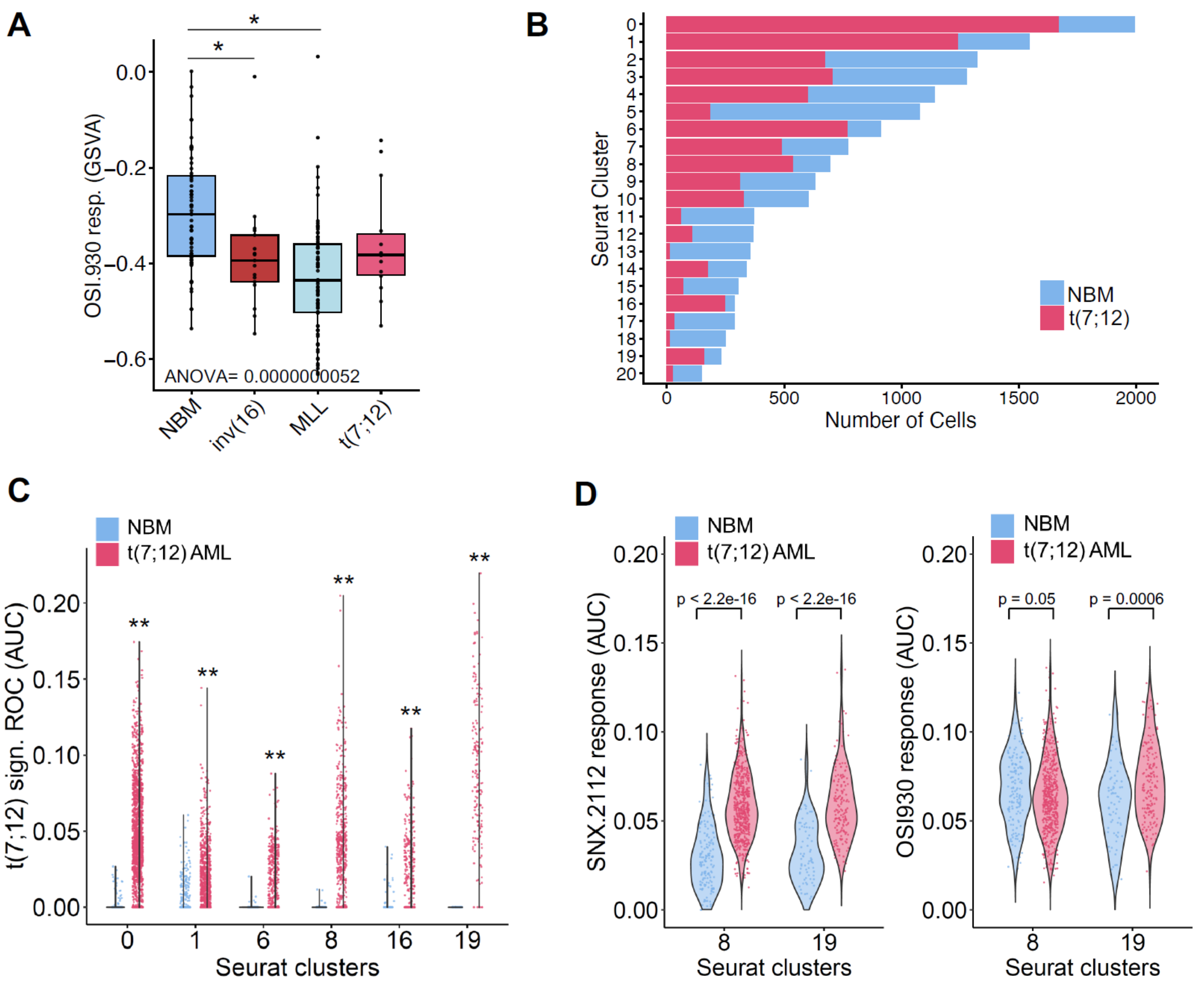


**<u>Figure 6 S1.</u> SNX.2112 selectively affects t(7;12)-AML signature t(7;12)-AML transcriptomes. A.** GSVA enrichment analysis of OSI.930 response signature in infant AML bulk-RNA seq from TARGET by subtype [inv(16), MLL/KMT2A, and t(7;12)] and normal bone marrow (NBM). Statistical testing was calculated by ANOVA and post-hoc pair-wise significance between classes by Tukey's test; p value ≤ 0.05 (*),0.001 (**), 0.0001 (***), and 0.00001 (****). **B.** Bar chart representing the proportion of cells populating each cluster by sample identity in scRNA-seq of t(7;12) AML vs normal bone marrow (NBM). **C.** Violin plot comparing AUC values for t(7;12) signature used for ROC analysis by cluster number and sample identity (normal bone marrow, NBM; t(7;12) AML patient), in clusters where t(7;12) cells are overrepresented over NBM. Statistical significance is determined by Wilcoxon test, p value ≤ 0.05 (*), 0.001 (**), 0.0001 (***), and 0.00001 (****). **D.** Violin plot comparing AUC values for SNX.2112 response (left) and OSI930 (right) by cluster number and sample identity (normal bone marrow, NBM; t(7;12) AML patient), in clusters where t(7;12) cells are overrepresented over NBM. Statistical significance is determined by Wilcoxon test, p value ≤ 0.05 (*), 0.001 (**), 0.0001 (***), and 0.00001 (****).

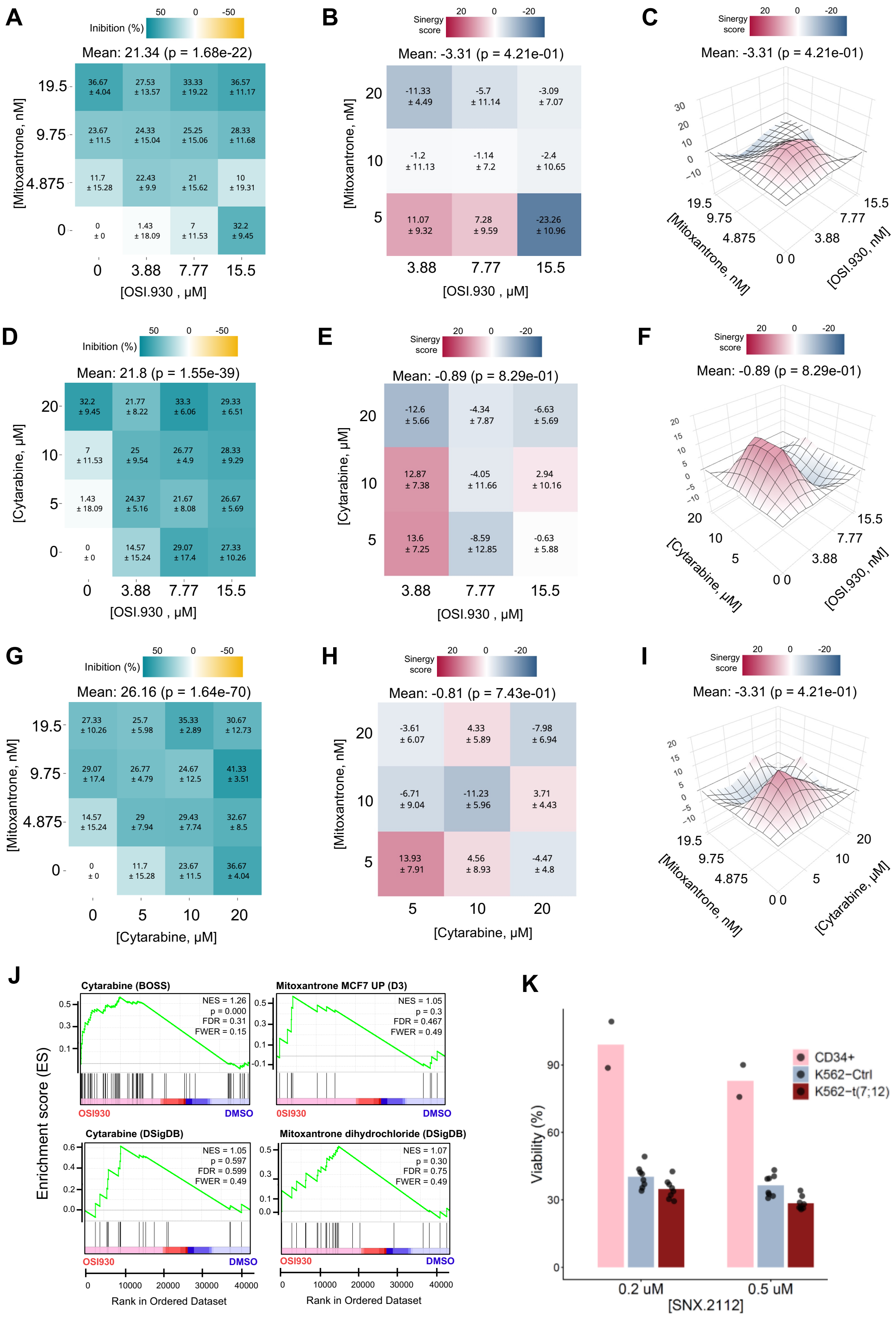


**Figure 7 S1. OSI.930 does not show synergistic effect in combination with chemotherapeutic agents. A-D.** Dose response map of K562-t(7;12) treated with OSI.930 in combination with mitoxantrone (**A.**) or with cytarabine (**D.**). Values of ¼ IC50, ½ IC50, and

IC50 of all drugs are shown. p values at the whole dose matrix level are shown at the top of the matrixes, together with the global mean. Individual scores for each combination are shown together with the ± value of the mean for each replicates set. **B-E.** HSA synergy heatmaps showing the synergy score of K562-t(7;12) treated with OSI.930 in combination with mitoxantrone (**B.**) or with cytarabine (**E.**). Values of ¼ IC50, ½ IC50, and IC50 of all the drugs are indicated. p values at the whole dose matrix level are shown at the top of the heatmaps, together with the global mean. Individual scores for each combination are shown together with the ± value of the mean for each replicates set. **C-F**. 3D representation of the **B.** and **E.** HSA synergy maps, respectively. **G.** Dose response map of K562-t(7;12) treated with cytarabine in combination with mitoxantrone. Values of ¼ IC50, ½ IC50, and IC50 of all the drugs are indicated. p values at the whole dose matrix level are shown at the top of the matrixes, together with the global mean. Individual scores for each combination are shown together with the ± value of the mean for each replicates set. **H-I.** HSA synergy heatmap (**H.**) or 3D map (**I.**) showing the synergy score of K562-t(7;12) treated with cytarabine in combination with mitoxantrone. Values of ¼ IC50, ½ IC50, and IC50 of all the drugs are indicated. p values at the whole dose matrix level are shown at the top of the matrixes, together with the global mean. Individual scores for each combination are shown together with the ± value of the mean for each replicates set. **J.** GSEA of the cytarabine (BOSS and DSigDB databases) and mitoxantrone (MCF7 UP(D3) and DSigDB databases) signatures against RNA-seq of K562-t(7;12) treated with OSI.930 (red) or DMSO (blue). **K.** Cell viability measured by MTT assay following SNX.2112 treatment for 24 hours. Viability is calculated on absorbance at 570 nm measured following incubation with MTT reagent relative to DMSO control treatment.
